# Neonatal sensory networks at birth predict cognitive, language, and motor outcomes at 18 months

**DOI:** 10.64898/2026.04.04.716445

**Authors:** Mi Zou, Arun L. W. Bokde

## Abstract

The relationship between neonatal brain activity patterns and later cognitive development has become a central topic in developmental neuroscience. Addressing this question requires whole-brain analytical approaches capable of identifying which large-scale functional systems carry stable and generalizable predictive signals. However, most existing studies remain focused on specific brain regions or localized functional circuits, such as thalamocortical pathways and amygdala-centered emotional networks. While these region-specific investigations have provided important insights, they are inherently limited in terms of robustness and cross-sample generalizability. As a result, systematic evidence identifying which large-scale functional systems reliably support stable and generalizable predictive signals remains scarce. Overcoming the methodological constraints of conventional whole-brain analytical paradigms has therefore become a key bottleneck in advancing our understanding of how early brain activity patterns relate to subsequent cognitive development.

Here, using data from 402 infants in the developing Human Connectome Project (278 term-born; 124 preterm-born), we introduce a region-of-interest (ROI)–constrained variant of Connectome-Based Predictive Modeling (CPM) that incorporates ROI-degree–guided feature selection to predict 18-month Bayley-III cognitive, language, and motor outcomes. Model performance declined as progressively lower-degree regions were included, indicating that conventional whole-connectome CPM may obscure robust predictive signals by incorporating low signal-to-noise (SNR) features.

Our models robustly predicted cognitive, language, and motor outcomes at 18 months of age. Cohort-specific connectivity patterns emerged. In term-born infants, dominant predictive features were concentrated in visual–auditory interactions, as well as connections between visual and auditory networks and other cortical regions. Interhemispheric and intrahemispheric connections contributed in roughly equal proportions. In contrast, among preterm infants, predictive features were primarily concentrated in connectivity involving auditory and temporoparietal networks, with interhemispheric connections comprising approximately twice the number of intrahemispheric connections. The whole-cohort model (term + preterm) reflected the combined contributions of both term- and preterm-associated connectivity patterns.

Predictions generalized across Bayley composite and subscale scores and were supported by permutation testing and held-out validation. These findings identify early sensory hubs—particularly visual and auditory regions—as promising early biomarkers for later neurodevelopmental outcomes. Furthermore, they demonstrate that ROI-constrained CPM can reveal meaningful predictive signals that may be obscured by conventional connectome-wide approaches.

## Introduction

Neonatal brain networks exhibit coherent functional organization from the earliest stages of life, with primary sensory and motor systems already identifiable in late gestation and at term-equivalent age(Doria et al., 2010; Fransson et al., 2009). This early topology scaffolds the emergence of later cognition, language, and motor function, positioning resting-state fMRI (rs-fMRI) as a promising source of biomarkers for neurodevelopment. Preterm birth (<37 weeks’ gestation) most commonly happens during the third trimester and is estimated to affect around 11% of all live births around the world(Chawanpaiboon et al., 2019). Recent neonatal studies have mapped modular, largely symmetric resting-state networks (RSNs) and documented preterm-related disruptions in connectivity, underscoring both the maturity of early sensory systems and the vulnerability of perinatal network development(Eyre et al., 2021). Understanding how early connectivity relates to later developmental outcomes, and how preterm birth may alter this relationship, is an important question to investigate, which may shed light on the development of early cognition and potentially offer early prevention methods for developmental delay.

Prior neonatal rs-fMRI studies have linked specific circuits (e.g., thalamocortical, DMN–ECN interplay, amygdala-centric networks) to later outcomes(Alcauter et al., 2014; Della Rosa et al., 2021; Rogers et al., 2017; Toulmin et al., 2021), and emerging work investigated the relationships between dynamic functional connectivity and later outcomes(França et al., 2024; Xu et al., 2024). However, most studies are constrained in at least one of three ways: (i) cohort scope (often preterm-only or small, clinically heterogeneous samples), (ii) network scope(specific circuits or brain regions); (iii) statistical framing (mainly association analysis), limiting generalizability to new individuals. Consequently, we have limited evidence about which large-scale systems carry robust, generalizable signal, and whether those systems differ between term- and preterm-born infants.

Here we address this gap using a large openly available dataset from the developing Human Connectome Project (dHCP). We developed an ROI-constrained variant of Connectome-Based Predictive Modeling to predict 18-month Bayley-III cognitive, language, and motor scores from neonatal rs-fMRI in term- and preterm-born infants (278 term; 124 preterm). Connectome-Based Predictive Modeling (CPM)(Shen et al., 2017), a data-driven framework that links functional connectivity patterns to behavioural outcomes, identifying edges most strongly associated with the target measure and builds predictive models from these connections. Hubs are highly interconnected and centrally positioned in the brain’s functional network, which makes their connectivity estimates more robust and their contributions to information processing more prominent (Power et al., 2013; van den Heuvel & Sporns, 2013). Our approach extended standard CPM by introducing a hub-oriented framework that isolates robust, biologically meaningful predictive signals. Specifically, we (i) implemented stability-driven ROI discovery, identifying and ranking regions of interest (ROIs) by their degree following extensive resampled edge selection, and (ii) applied ROI-constrained feature selection, restricting predictive features to edges connected to the high-degree ROIs. This design enhanced model interpretability and reliability by focusing on network hubs with intrinsically higher SNR. Empirically, prediction accuracy declined as lower-degree regions were added, consistent with a reduction in SNR. Permutation testing on the mean predictive correlation across ROI-set sizes confirmed that the observed performance exceeded chance levels. Thus, our hub-constrained CPM uncovers meaningful connectivity–behavior relationships that are often obscured in conventional CPM by the inclusion of low-connectivity regions.

We report three main findings. First, neonatal connectivity in primary sensory systems—particularly visual and auditory cortices—predicts subsequent cognitive, language, and motor outcomes, consistent with the early maturation of these networks. Second, predictive anatomy differs by cohort: in term infants, prediction emphasizes visual–auditory network interaction, whereas in preterm infants it emphasizes temporoparietal and auditory networks with a marked interhemispheric bias. Third, predictive models in the whole cohort reflect contributions from both term- and preterm-associated connectivity patterns, indicating convergent pathways by which early networks relate to later behaviour. Methodologically, we introduce a ROI-constrained CPM that reveals stable and interpretable connectivity–behavior relationships that are often masked in conventional CPM by noise from low-connectivity regions.

## Methods

### Participants

We analyzed data from 402 participants selected from the 814 available functional MRI (fMRI) datasets in the Developing Human Connectome Project(dHCP, Release 4). Among 124 preterm-born infants, 91 had two imaging sessions; in such cases, only the first session, scanned shortly after birth, was included in the analysis. Participants were excluded based on the following criteria: (i) excessive head motion (n = 151; motion exclusion criteria see below), (ii) missing Bayley-III Composite scores at 18 months (n = 116), and (iii) term-born infants with postnatal age (scan age - gestational age) >3 weeks (n = 54). The final cohort comprised 278 term-born and 124 preterm-born infants, with preterm defined as gestational age <37 weeks. Demographic information is presented in Table 1. Neurodevelopmental outcomes were assessed at 18 months corrected age using the Bayley Scales of Infant and Toddler Development, Third Edition(Bayley-III).

**Table 1.** Research participants.

|  | Term(n=278) | Preterm(n=124) | Statistics | *p* value |
| --- | --- | --- | --- | --- |
| Demographics |  |  |  |  |
| GA at birth,<br>median(range),weeks | 40.14 (37-42.7) | 33.36 (23.0-36.9) | 16.0<sup>a</sup> | p < 0.001 |
| PMA at scan,<br>median(range),weeks | 41.07 (37.4-44.7) | 35.43 (27.3-45.1) | 13.8<sup>a</sup> | p < 0.001 |
| Female(%) | 126 (45) | 53 (43) | 0.2<sup>b</sup> | 0.631 |
| Weight at birth,<br>median(range),kg | 3.77 (2.7-4.3) | 1.64 (0.5-4.1) | 10.9<sup>a</sup> | p < 0.001 |
| % FD outliers(SD) | 4.64 (2.7) | 4.65 (2.5) | -0.2<sup>a</sup> | p = 0.854 |
| Follow-up |  |  |  |  |
| motor composite standardized<br>score, mean(SD)<sup>c</sup> | 102.06 (9.9) | 99.71 (11.2) | 1.7<sup>a</sup> | p = 0.086 |
| gross<sup>c</sup> | 9.25 (1.8) | 8.94 (1.9) | 1.7<sup>a</sup> | p = 0.087 |
| fine<sup>c</sup> | 11.37 (2.3) | 10.88 (2.5) | 1.5<sup>a</sup> | p = 0.133 |
| Language composite<br>standardized score, mean(SD)<sup>c</sup> | 98.11 (16.0) | 96.34 (17.1) | 0.4<sup>a</sup> | p = 0.670 |
| expressive<sup>c</sup> | 9.10 (2.7) | 8.52 (2.7) | 1.8<sup>a</sup> | p = 0.067 |
| receptive<sup>c</sup> | 10.19 (3.3) | 10.17 (3.6) | -0.7<sup>a</sup> | p = 0.508 |
| Cognitive composite<br>standardized score, mean(SD)<sup>c</sup> | 101.04 (11.2) | 98.67 (13.2) | 1.5<sup>a</sup> | p = 0.135 |

FD framewise displacement <sup>a</sup>Z(Mann-Whitney U test) <sup>b</sup> $\chi^2$ -test <sup>c</sup>Bayley-III Scales of Infant Development (Third Edition)

### MRI Data Acquisition

Functional MRI data were collected as part of the developing Human Connectome Project (dHCP) at the Evelina Newborn Imaging Centre, Evelina London Children’s Hospital, utilizing a 3 Tesla Philips Achieva system. The study received ethical clearance from the UK National Research Ethics Authority (14/LO/1169), and all participating families provided written informed consent prior to the imaging sessions. All scans were performed without the use of sedation in a neonatal environment designed specifically for safety and comfort, which included a custom 32-channel head coil, an acoustic hood, and devices to ensure proper positioning of the infants. Infants wore MRI-compatible ear protection to reduce noise exposure, and a neonatal nurse or paediatrician continuously monitored vital signs, including heart rate, oxygen saturation, and body temperature. Blood-oxygen-level-dependent (BOLD) fMRI data were acquired using a multi-slice echo planar imaging sequence with multiband excitation (factor 9) over a scan duration of 15 minutes and 3 seconds, producing 2300 volumes. Imaging parameters included a repetition time (TR) of 392 ms, echo time (TE) of 38 ms, a flip angle of 34°, and a voxel resolution of 2.15 mm isotropic. High-resolution T2w anatomical images were obtained for structural analysis and functional data registration. T2-weighted images were obtained with a resolution of 0.8 mm isotropic and a field of view of 145 × 145 × 108 mm, a TR of 12 s, and a TE of 156 ms.

### Functional data preprocessing

The preprocessing of neuroimaging data was performed using a bespoke pipeline specifically optimized for neonatal imaging and developed for the dHCP, as detailed in Fitzgibbon et al. (2020). The pipeline accounted for susceptibility-induced dynamic distortions as well as intra-and inter-volume motion artifacts. Twenty-four extended rigid-body motion parameters were regressed alongside single-subject independent component analysis (ICA) noise components identified using the FSL FIX tool (Oxford Centre for Functional Magnetic Resonance Imaging of the Brain’s Software Library, version 5.0). The denoised data were first registered to the T2-weighted native space using boundary-based registration (Greve & Fischl, 2009). They were then non-linearly transformed to a standard space with a weekly template from the dHCP volumetric atlas through diffeomorphic multimodal (T1/T2) registration (Avants et al., 2008).

Because head motion is a potential surrogate marker for the infant’s arousal state and can interact with underlying neural activity, we adopted a conservative approach to minimize the impact of motion-related artifacts. Specifically, for each subject, we selected a continuous subset (1600 from the original 2300 acquired volumes) with the minimum total framewise displacement (FD) and the dataset was cropped accordingly – the cropped subset was used for all subsequent analyses. After the cropping, volumes were flagged as motion outliers based on FD, with outliers identified as FD >1.5 interquartile range (IQR) above the 75th centile. Subjects with more than 160 motion-outlier volumes (>10% of data) were excluded from further analyses. The motion outliers of the resulting data from the participants are showed in Table 1. No significant differences were observed between term and preterm groups under the assumption of non-normality (p = 0.854, Mann-Whitney U-test). The number of outliers was included as a covariate in subsequent regression analyses.

### Connectivity Matrices

Whole-brain functional connectivity was computed using the CONN toolbox (Whitfield-Gabrieli & Nieto-Castanon, 2012). Brain regions were defined using the Schaefer 200-node cortical atlas(Schaefer et al., 2018) combined with 8 subcortical regions from the dHCP parcellation, yielding 208 nodes. The atlas was aligned to the 40-week template via rigid-body, affine, and SyN diffeomorphic transformations implemented in ANTs(Avants et al., 2008). Mean time courses were extracted for each ROI, and pairwise Pearson correlations were computed and Fisher Z-transformed, resulting in a symmetric 208×208 connectivity matrix per subject.

### ROI-constrained Connectome–Based Predictive Modeling (R-CPM)

The pipeline was applied separately to preterm-only, term-only, and whole cohort (term + preterm), with preterm status included as a covariate in the whole cohort analysis. All analyses are performed separately for positive and negative associations.

Stage 1 — Stable edge discovery: We ran 10-fold repeated 150 times with new partitions. For each fold we computed partial correlations between each edge and behaviour controlling for gender, head motion (number of outliers), age at scan, and preterm status when applicable .

Edges with r>0 and p<0.05 were counted as positive; r<0 and p<0.05 as negative. Counts were accumulated across all folds and repeats. An edge was labelled stable if selected in ≥95% of the 10×150 iterations. ROI ranking(below) was performed separately for positive and negative graphs.

Stage 2 — ROI ranking: From the Stage-1 stable graph (sign-specific), we identified hubs by iteratively removing the highest-degree ROI and recomputing degrees on the remaining graph. The ROI with the largest degree was labelled the top hub; after removing it and its incident edges, the ROI with the largest updated degree was labelled second, and so on, yielding a ROI order 𝑅 = (𝑟_1_, 𝑟_2_, …) (Figure 1a). No Stage-1 edges were carried forward as features; only the ROI order 𝑅 was passed to Stage 3.

**Figure 1.**
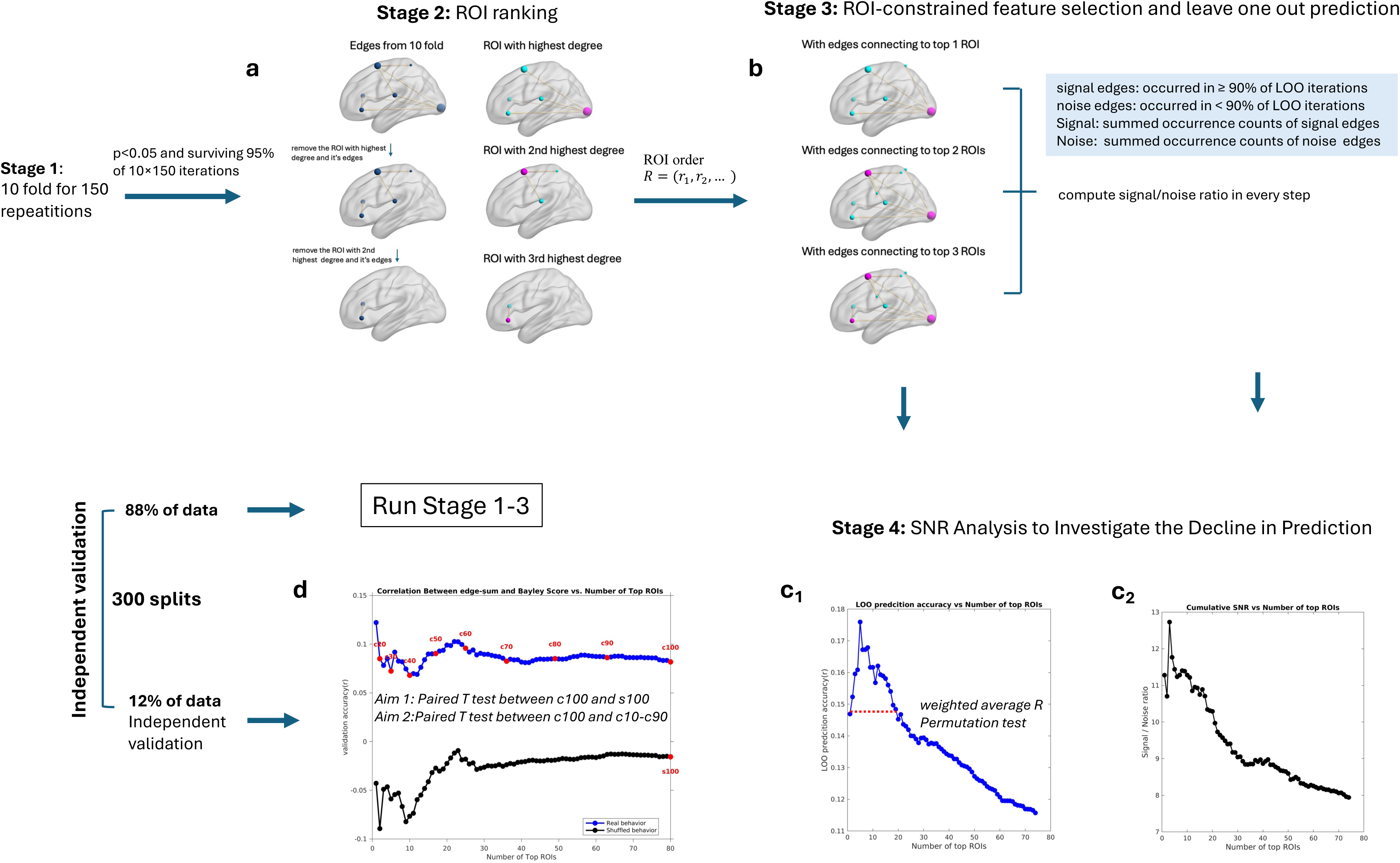
ROI-constrained CPM pipeline with ROI-constrained feature selection and independent validation. (a) Stage-2 — ROI ranking: ROIs were sorted by iteratively removing the highest-degree ROI to define a high-to-low ROI degree order. No Stage-1 edges were carried forward as features; only the ROI order 𝑅 was passed to Stage 3. (b) Stage-3 — ROI-constrained CPM : For each ROI-set size (n = 1…|R|), edges that survived p<0.05 and connected to at least one of the top-n ROIs were selected via LOO to calculate prediction accuracy. SNR was also computed for each prefix n by dividing the summed occurrence counts of signal edges (occurred in ≥ 90% of LOO iterations) by those of noise edges (occurred in ≥ 90% of LOO iterations);(c) Stage-4 — SNR analysis: As progressively lower-degree ROIs were added, SNR steadily decreased(c_2_), mirroring the decline in predictive accuracy(c_1_). And SNR and prediction accuracy were strongly correlated (r = 0.96, p < 0.0001). We conducted a permutation test to confirm that the observed weighted average prediction performance (weighted by degree of each ROI) across all ROI-set sizes n exceeded chance; (d) Independent validation: For each ROI-set size (n = 1…|R|), validation subjects’ network strength—sums of signal edges incident on the top-n ROIs—were correlated with behavior to yield 𝑟(n) curves. The cumulative distribution of edge counts across ROIs was computed, and thresholds were set at 10 %, 20 %, …, 100 % coverage of total edges. Aim 1 — Check whether all signal edges contribute predictive information: Paired T test between observed and shuffled behavior at coverage 100%. Aim 2 — Check whether the predictive effect declines as signal edges of low degree ROIs are added: Paired t-tests compared performance at each intermediate coverage level (10–90 %) with that at 100 %. Abbreviations: c10, coverage 10%; s100, shuffled behavior at coverage 100%; ROI, region of interest; LOO, leave one out; SNR, signal to noise ratio.

Stage 3 — ROI-constrained feature selection and leave-one-out (LOO) prediction: We adopted a ROI-constrained variant of CPM. For each prefix size 𝑛 = 1, …, ∣ 𝑅 ∣(looping over the ordered ROIs) and each sign, we performed LOO: (1) Feature selection: Within each LOO training set (excluding the held-out subject), we computed partial correlation between each edge and behaviour(the same covariates applied), retaining edges meeting p<0.05 that connected to at least one of the top-𝑛 ROIs (Figure 1b). (2) Summary edges: For each subject computed 𝑆^+^(sum of positive selected edges) and 𝑆^−^(sum of negative selected edges). (3) Prediction: We fitted separate linear models on the LOO training set for 𝑆^+^and 𝑆^−^and predicted behaviour of the left-out subject. (4) Edge stability across LOO: After LOO completed for a given 𝑛, retaining edges occurring in ≥90% of LOO iterations as 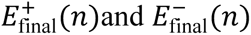. Stage 3 yielded, for each 𝑛, sign-specific LOO accuracy (Figure 1c_1_) and training-stable edge sets. As figure 1c_1_ shows, as the number of top ROIs increases, the prediction accuracy goes down.

Stage 4 —Signal to noise ratio (SNR) analysis to investigate the decline in prediction: To investigate why the predictive accuracy decreased as more ROIs were added, we examined how the inclusion of lower-degree ROIs affected the SNR.

For each prefix size *n* = 1,…,|R| (looping over the ordered ROIs), we recorded occurrence of all edges across the LOO iterations in Stage 3. These edges were divided into two categories based on a stability threshold of 0.9 × (number of LOO iterations): (1) Signal edges: edges with occurrence counts ≥ threshold; (2) Noise edges: edges with occurrence counts < threshold.

The SNR for each prefix *n* was then computed as

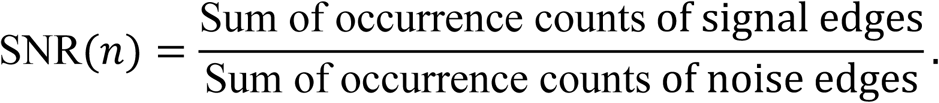

As progressively lower-degree ROIs were added in the whole-cohort model of composite cognition, the SNR consistently declined (Figure 1c_2_). And SNR was strongly correlated with LOO prediction accuracy (r = 0.96, p < 0.0001; Figures 1c_1_–1c_2_). Similar relationships between SNR and prediction accuracy were observed across all behaviours and cohorts (Supplementary Table 1), indicating that the decrease in prediction arises from a reduced SNR as low degree ROIs are incorporated.

### Permutation testing of prediction accuracy

After demonstrating that the decline in predictive accuracy with increasing number of top ROIs reflects a reduction in SNR, we conducted a permutation test to confirm that the observed weighted average prediction performance (weighted by degree of each ROI) across all ROI-set sizes *n* (figure 1c_1_) exceeded chance. Across Stages 1–3, behavioural labels were randomly permuted while preserving the full analytical pipeline. Because Bayley scores in this healthy neonatal cohort included many tied values, we excluded permutations in which >40% of the scores were reassigned to their original positions. In cases where no edges survived across the 10 folds in Stage 1, the corresponding prediction performance was assigned a value of zero. For each permutation, we computed the weighted average R across all ROI-set sizes *n*. The one-sided *p*-value was calculated as following:

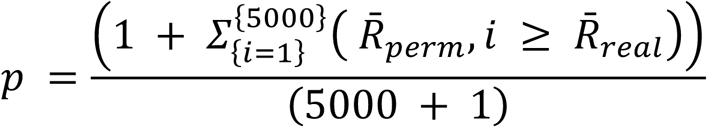

5000= total number of permutations. 𝑅̄_perm,i_ = weighted average correlation between predicted and permuted behavioral scores across all ROI set sizes *n*. 𝑅̄_real_= weighted average correlation between predicted and observed behavioral scores across all ROI set sizes *n*.

### Independent validation of signal edge effects and ROI-dependent decline

We had previously established that the reduction in predictive accuracy is driven by the growing contribution of noise edges originating from low-degree ROIs. It remains unclear whether all signal edges—including those connected to low-degree ROIs—contribute predictive information, and whether prediction accuracy decreases as signal edges from low-degree ROIs are added.

In order to investigate this, we conducted an independent validation analysis. Approximately 88% of the data were used for running stage 1-3, while the remaining 12% were reserved for independent validation. The split was repeated for 300 times. All feature discovery and model fitting used training data only; validation data were untouched until final testing.

To preserve the discretized outcome distribution, we stratified the split by exact Bayley-III score value: for values with ≥8 subjects we assigned ∼88% to training and ∼12% to validation; values with <8 subjects were pooled before applying the same proportion.

For each sign and each prefix size *n = 1,…,|R|* (looping over the ordered ROIs), we used the training-stable edge sets 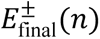(signal edges from Stage 3): (1)For each validation subject, we computed 𝑆^+^/ 𝑆^−^ by summing all edges incident on the top-*n* ROIs; (2)Each subject’s 𝑆^±^ was correlated (Pearson’s *r*) with behaviour to yield predictive-accuracy curves 𝑟^±^(𝑛); (3) A single-shuffle was performed by permuting the behaviour once per split to estimate chance-level 𝑟^±^(𝑛)values (Figure 1d).

Aim 1 — Do all signal edges contribute predictive information?

To evaluate whether the entire set of signal edges retains predictive power, we focused on the largest ROI set (*n = |R|*) and did paired *t*-tests(fisher-z transformed r value) between real behaviour and shuffled behaviour across 300 splits to test whether the final model’s correlation remained significantly above the single shuffle (Figure 1d).

Aim 2 — Does the predictive effect decline as signal edges of low degree ROIs are added? To examine how expanding the ROI set influences performance, we quantified accuracy as a function of cumulative edge coverage. For each behavior, the cumulative distribution of edge counts across ROIs was computed, and thresholds were set at 10 %, 20 %, …, 100 % of total edges. For each coverage level, the corresponding predictive correlations 𝑟(𝑛) were extracted. Across 300 splits, paired *t*-tests (Fisher-z transformed r value) compared performance at each intermediate coverage level (10–90 %) with that at 100 % (Figure 1d).

### Handling Clustered Behavioural Scores in Term-Cohort Validation

The Bayley-III scores of the term cohort were even more clustered and contained repeated values. To improve SNR in validation, we fit a covariate-only linear model within training (gender, motion, age; plus preterm only in whole-cohort analyses) and get covariate coefficients. Then we applied the covariate coefficients derived from the training to the validation subjects to obtain residualized validation behaviour. Validation accuracy was calculated by the correlation between sum of edges 𝑆^±^ and residualized behaviour.

The permutation p-value of average R across all ROI-set sizes was not significant in term cohort(see Table 2). To further demonstrate the effect in term cohort, we performed a permutation test that incorporated the independent validation procedure (1,000 iterations). To avoid potential confounding from covariates, the original behavioural scores were used instead of residualized scores in the independent validation. For each permutation, the independent validation was repeated across 200 random splits to obtain a mean validation correlation(r̄_validation_). A one-sided p-value for the observed mean validation correlation was then computed based on the 1,000 permuted mean validation correlations.

**Table 2.** Statistics of permutation and independent validation.

|  | Permutation |  |  |  | Independent validation |  |  |  |  |  |
| --- | --- | --- | --- | --- | --- | --- | --- | --- | --- | --- |
| | weighted average R<sup>a</sup> | | p<sup>a</sup> | | $\bar{r}_{\text{validation}}^{\text{c}}$ | 95% CI<sup>c</sup> | $\bar{r}_{\text{shuffled}}^{\text{c}}$ | t(299)<sup>c</sup> | p<sup>c</sup> | Cohen's $d_z^{\text{c}}$ |
| whole cohort (n=402) |  |  |  |  |  |  |  |  |  |  |
| composite cognition | 0.15 |  | 0.019 |  | 0.12 | 0.11-0.14 | -0.009 | 11.8 | 1.75E-26 | 0.68 |
| composite language | 0.15 |  | 0.015 |  | 0.13 | 0.11-0.14 | 0.003 | 10.9 | 2.02E-23 | 0.63 |
| expressive language | 0.17 |  | 0.008 |  | 0.16 | 0.14-0.17 | 0.008 | 12.4 | 6.20E-29 | 0.72 |
| receptive language | 0.12 |  | 0.072 |  | 0.09 | 0.08-0.11 | -0.019 | 8.0 | 3.57E-14 | 0.46 |
| composite motor | 0.13 |  | 0.056 |  | 0.08 | 0.06-0.09 | -0.008 | 6.4 | 7.53E-10 | 0.37 |
| fine motor | 0.13 |  | 0.042 |  | 0.06 | 0.04-0.07 | -0.007 | 3.2 | 0.002 | 0.18 |
| gross motor | 0.11 |  | 0.079 |  | 0.07 | 0.05-0.08 | -0.008 | 5.2 | 4.09E-07 | 0.30 |
| preterm cohort (n=124) |  |  |  |  |  |  |  |  |  |  |
| composite cognition | 0.28 |  | 0.004 |  | 0.12 | 0.09-0.14 | 0.020 | 5.1 | 4.95E-07 | 0.30 |
| composite language | 0.25 |  | 0.032 |  | 0.09 | 0.06-0.12 | -0.024 | 4.1 | 5.33E-05 | 0.22 |
| receptive language | 0.27 |  | 0.009 |  | 0.08 | 0.05-0.12 | -0.002 | 4.0 | 7.59E-05 | 0.23 |
| composite motor | 0.20 |  | 0.075 |  | 0.10 | 0.07-0.13 | 0.003 | 4.2 | 3.76E-05 | 0.24 |
| gross motor | 0.22 |  | 0.042 |  | 0.09 | 0.07-0.11 | 0.006 | 5.4 | 1.37E-07 | 0.31 |
| term cohort (n=278)<br>(negative network) |  |  |  |  |  |  |  |  |  |  |
| receptive language | 0.17 |  | 0.031 |  | -0.09 | -0.11- -0.08 | 0.004 | -6.0 | 5.54E-09 | -0.35 |
| gross motor | 0.15 |  | 0.058 |  | -0.07 | -0.1- -0.06 | 0.002 | -5.3 | 1.87E-07 | -0.31 |
|  | further permutation |  |  |  |  |  |  |  |  |  |
| term cohort (n=278) | weighted<br>average R<sup>a</sup> | p<sup>a</sup> | $\bar{r}_{\text{validation}}^{\text{b}}$ | p<sup>b</sup> | | | | | | |
| composite cognition | 0.10 | 0.374 | 0.04 | 0.009 | 0.10 | 0.08-0.11 | 0.007 | 6.6 | 1.50E-10 | 0.38 |
| composite language | 0.11 | 0.343 | 0.05 | 0.005 | 0.14 | 0.12-0.16 | 0.013 | 9.9 | 3.87E-20 | 0.57 |
| expressive language | 0.14 | 0.189 | 0.07 | 0.002 | 0.16 | 0.15-0.18 | 0.002 | 11.9 | 5.79E-27 | 0.69 |
| receptive language | 0.07 | 0.452 | 0.03 | 0.011 | 0.07 | 0.05-0.09 | 0.001 | 5.3 | 2.84E-07 | 0.30 |
| composite motor | 0.09 | 0.412 | 0.02 | 0.023 | 0.05 | 0.03-0.06 | 0.012 | 2.8 | 0.005 | 0.16 |
| fine motor | 0.10 | 0.391 | 0.02 | 0.022 | 0.05 | 0.03-0.06 | 0.003 | 3.3 | 0.001 | 0.19 |

<sup>a</sup>Permutation test: We conducted a permutation test (5000 iterations) to confirm that the observed weighted average prediction performance (weighted by degree of each ROI) across all ROI-set sizes (Figure 1c1) exceeded chance levels. Across Stages 1–3 (Figure 1), behavioral labels were randomly permuted while maintaining the full analytical pipeline. For each permutation, the average correlation R across all ROI-set sizes was computed. A one-sided p-value was then computed as the proportion of permutations whose weighted average correlation exceeded the observed value.
<sup>b</sup>For the term group, we conducted an additional permutation test that incorporated the independent validation procedure (1000 iterations). For each permutation, the full validation workflow was repeated for 200 random train–validation splits, yielding a mean validation correlation ( $\bar{r}_{\text{val\_per}}$ ). The one-sided p-value was then computed as the proportion of permutations in which $\bar{r}_{\text{val\_per}}$ exceeded the observed mean validation correlation.
<sup>c</sup>Independent validation: Significance of validation versus the behavior-shuffle null on the largest ROI set ( $n = |R|$ ) is tested with a paired t-test across 300 splits; we report $t(df)$ , exact $p$ , Cohen's $d_z$ , pooled $\bar{r}_{\text{validation}}$ with 95% CI and $\bar{r}_{\text{shuffled}}$ . Note: The difference between $\bar{r}_{\text{validation}}^b$ and $\bar{r}_{\text{validation}}^c$ arises because $\bar{r}_{\text{validation}}^c$ was computed using covariate-residualized behaviour in the independent validation, whereas $\bar{r}_{\text{validation}}^b$ used the observed (non-residualized) behaviour.

### Statistical evaluation and reporting

The following two statistical test significance will be reported: (1) permutation test of average R across all ROI-set sizes *n*; (2) independent validation test on the largest ROI set (*n = |R|*). Pooled mean validation accuracy (r̄) at n = R with 95% confidence interval will be reported.

Significance versus the behaviour-shuffle null at 𝑛 = 𝑅 is tested with a paired t-test across splits; we report 𝑡(df), exact 𝑝, and Cohen’s 𝑑_3_(see Table 2). Correlations were Fisher-z transformed for inference and back-transformed for presentation.

We considered two cases. First, when performance at 100% coverage was significantly above chance, we further asked whether 100% coverage also yielded the best validation accuracy compared to other coverages. In most behaviors, the highest validation accuracy was observed at 100% coverage, indicating that predictive performance generally improved as signal edges associated with low-degree ROIs were added. In this case, we reported the signal edges that survived in 90% of splits at 100% coverage. Second, for behaviors in which the best coverage was below 100%, this was typically because a small subset of top ROIs accounted for a disproportionately large share of the total edges (see Supplementary Tables 2–4 and Supplementary Figures S1, S5, and S9). If 100 % coverage is still significant above chance, we still reported the signal edges that survived in 90% of splits at 100% coverage(we compared the connectivity patterns at the best coverage and at 100% coverage and found them to be consistent,see Supplementary Figures S2–4, S6–8, and S10–15). If performance at 100% coverage was not significantly above chance, we report the signal edges that survived in 90% of splits at the last coverage level that remained significantly above chance. In both cases, all other tested coverage levels before the coverage we report also showed significant differences between real and shuffled data (paired t-test, p < 0.05).

The connectivity results are presented in a network format. Resting-state networks were defined using group-level independent component analysis of term-born infants scanned at 43.5–44.5 weeks postmenstrual age in the developing Human Connectome Project (dHCP) dataset (Eyre et al., 2021). Each ROI from the parcellation was assigned to the network with the highest spatial overlap (winner-takes-all criterion). The network map is shown in supplementary Figure S16.

## Results

In summary, visual and auditory cortices were core regions that contribute to prediction of cognition, language and motor Bayley score at 18 months of age in the whole (term + preterm neonates) group and term group, particularly notable was the connectivity between visual and auditory cortices. In contrast, preterm infant cohort, the prediction models relied more on auditory and temporoparietal networks. Results that passed at least one of the two statistical tests—the permutation test or the independent validation—are summarized in Table 2, and the corresponding connectivity patterns are illustrated in Figures 2–7.

**Figure 2.**
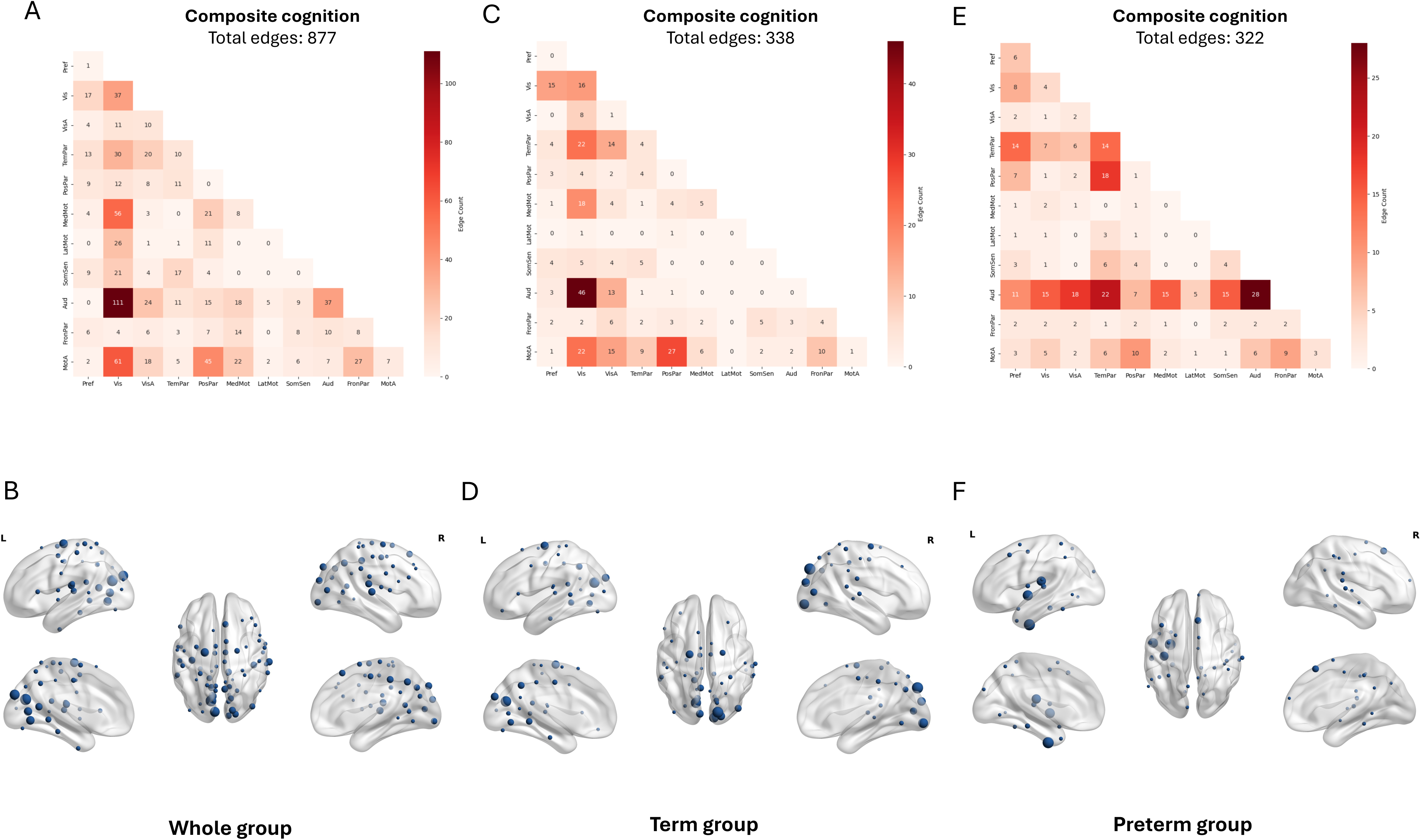
Cohort-specific predictive networks for cognition. (A,C,E) Connections plotted as the number of edges within and between each pair of canonical networks for the whole cohort, term-born cohort, and preterm-born cohort, respectively(left to right). (B,D,F) High-degree ROIs for the same models. Only ROIs with a degree ≥ one-sixth of the highest ROI are displayed; node size is proportional to degree. Abbreviations: Pref: prefrontal network; Vis: visual network; VisA: visual assosciation network; TemPar: temporoparietal network; PosPar: postparietal network; MedMot: medial motor network; LatMot: lateral motor network; SomSen: somotasesory network; Aud: auditory network; FronPar: Frontal Parietal network; MotA: motor assosciation network.

**Figure 3.**
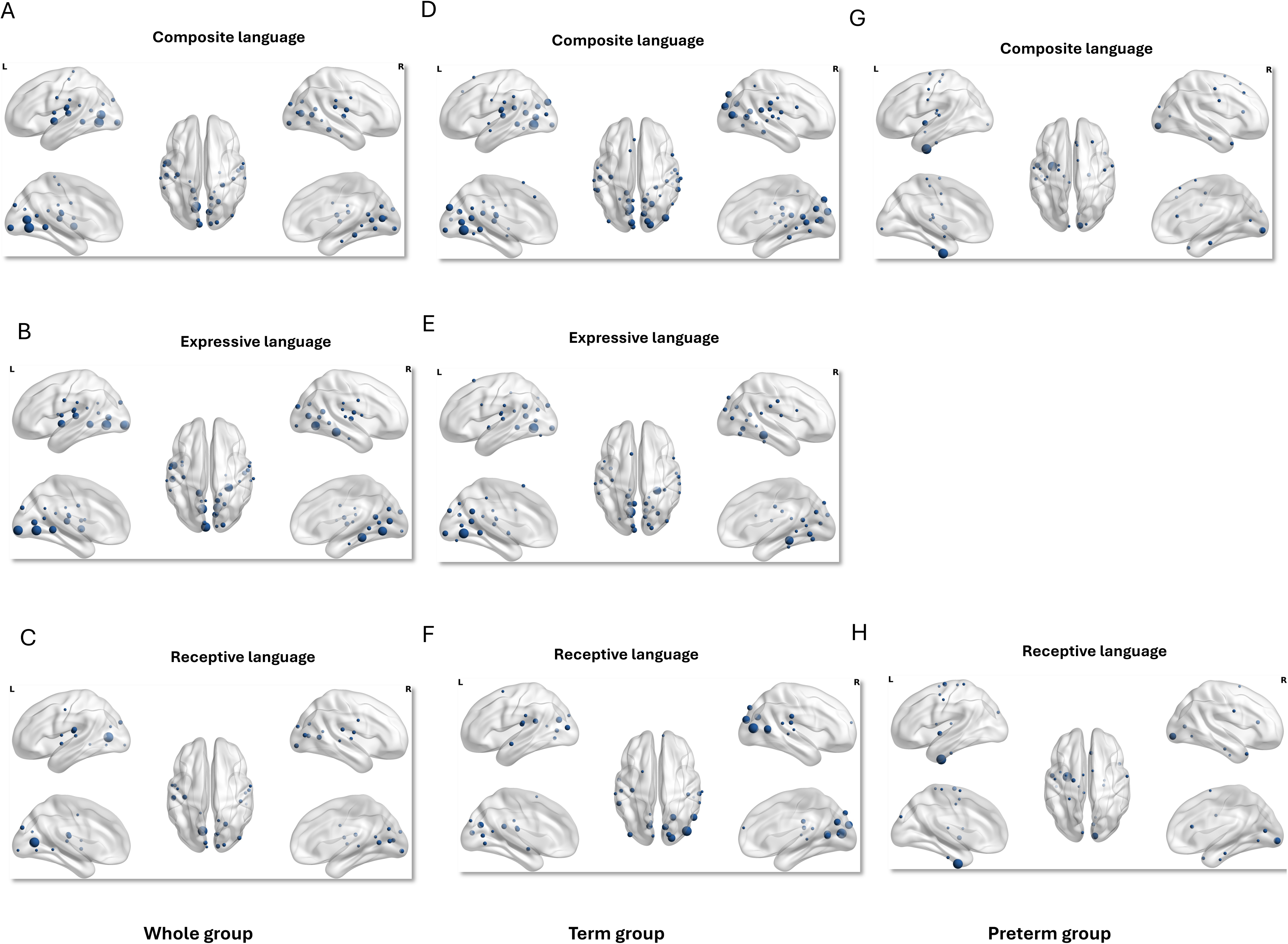
Predictive ROIs for language: high-degree regions across cohorts and subscales. High-connectivity regions are presented for the whole cohort, term-born cohort, and preterm-born cohort (left to right) across composite, expressive, and receptive language scores (top to bottom). Only ROIs with a degree ≥ one-sixth of the highest ROI are displayed; node size is proportional to degree.

**Figure 4.**
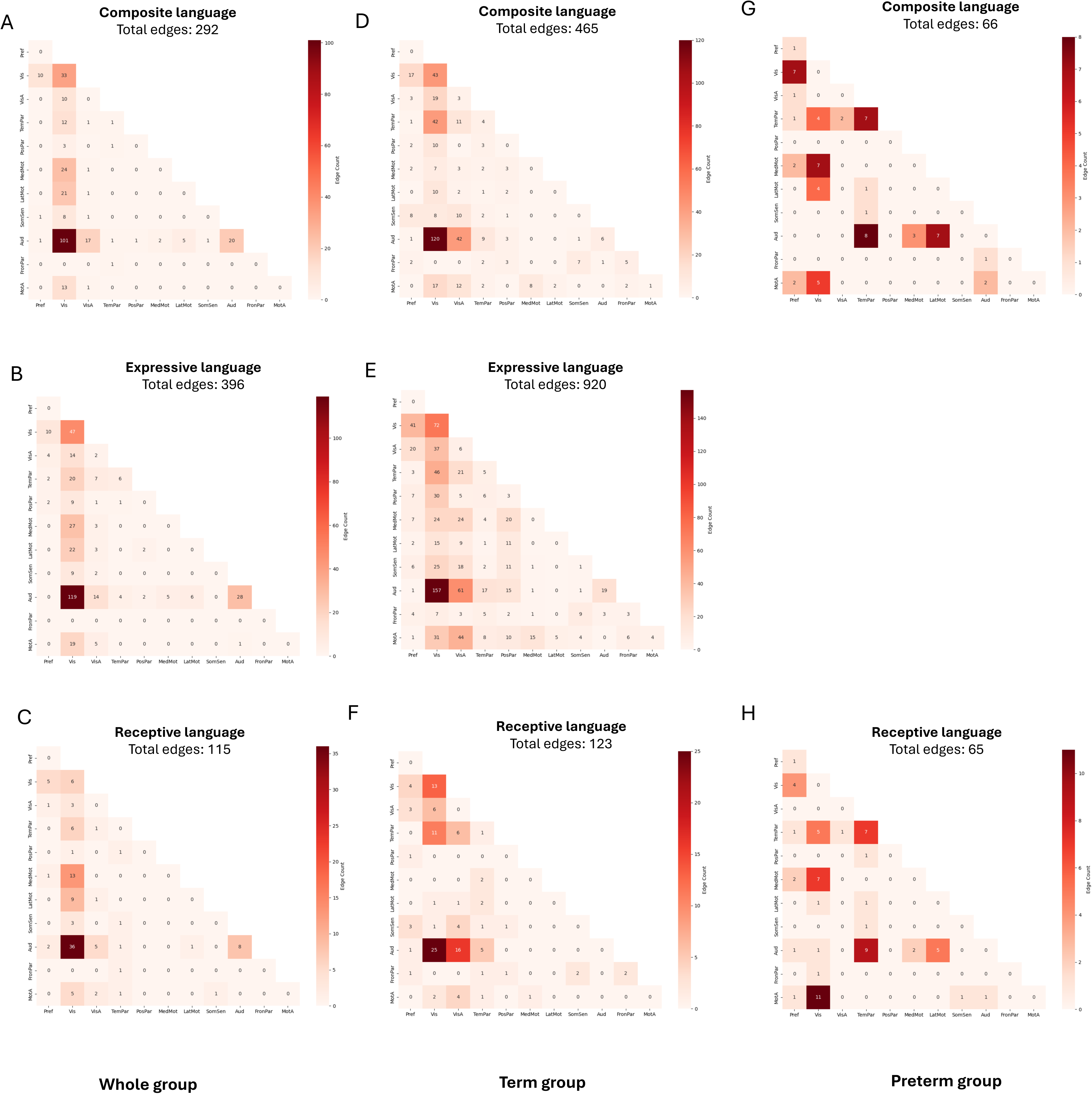
Predictive networks for language across cohorts and subscales. Connections plotted as the number of edges within and between each pair of canonical networks for the whole cohort, term-born cohort, and preterm-born cohort respectively(left to right) across composite, expressive, and receptive language scores (top to bottom).

**Figure 5.**
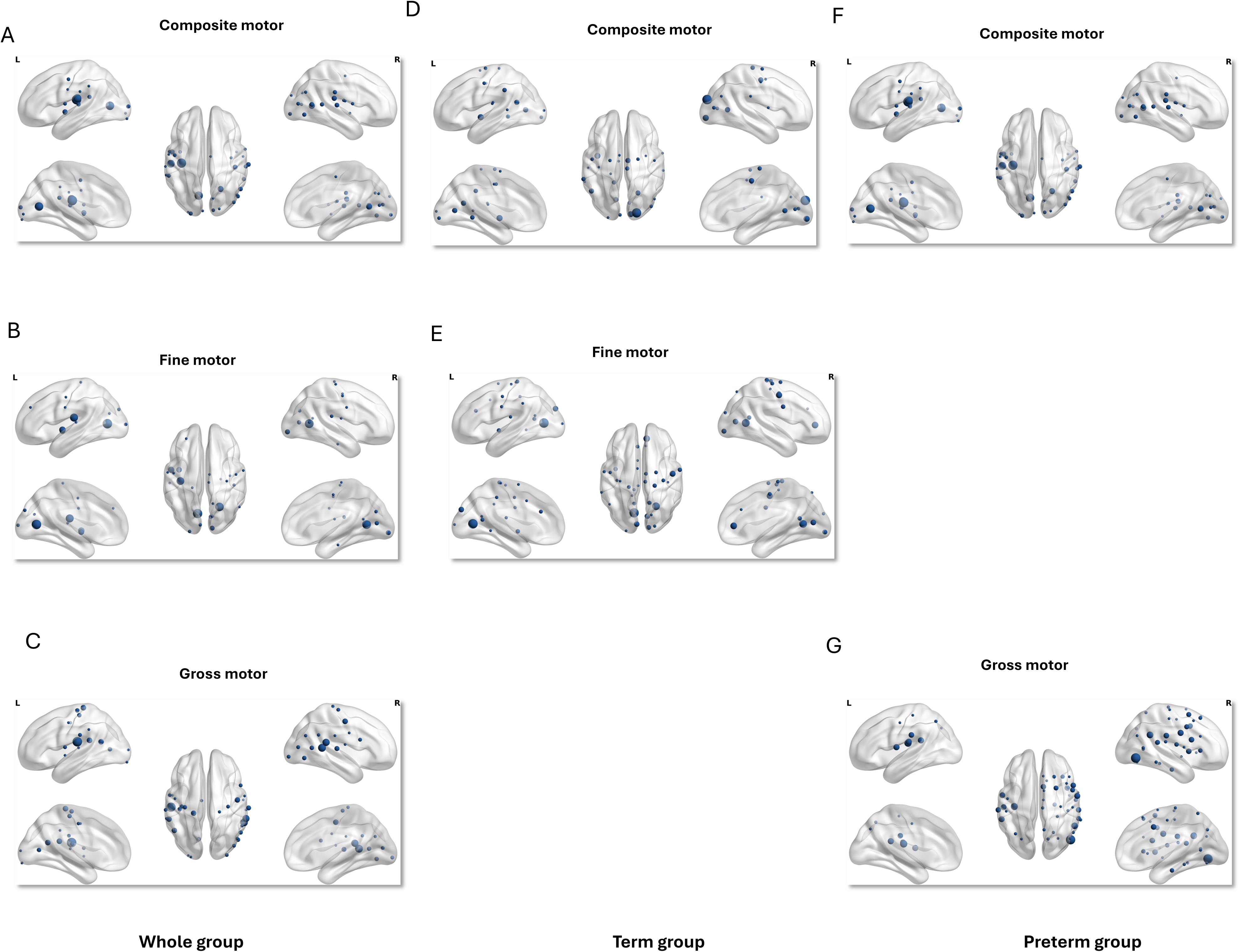
Predictive ROIs for motor: high-degree regions across cohorts and subscales. High-connectivity regions are presented for the whole cohort, term-born cohort, and preterm-born cohort (left to right) across composite, fine, and gross motor scores (top to bottom). Only ROIs with a degree ≥ one-sixth of the highest ROI are displayed; node size is proportional to degree.

**Figure 6.**
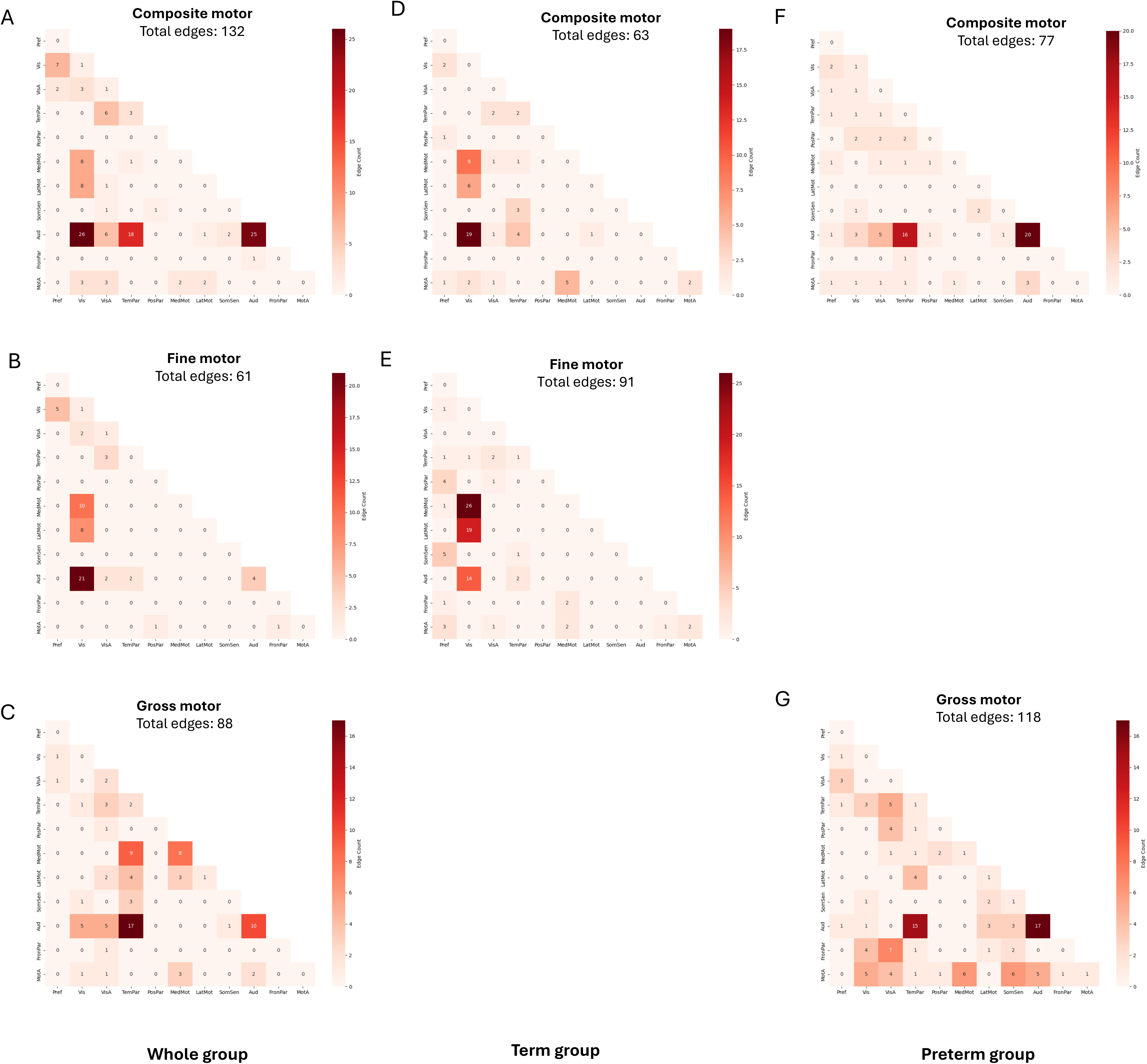
Predictive networks for motor across cohorts and subscales. Connections plotted as the number of edges within and between each pair of canonical networks for the whole cohort, term-born cohort, and preterm-born cohort (left to right) across composite, fine, and gross motor scores (top to bottom).

**Figure 7.**
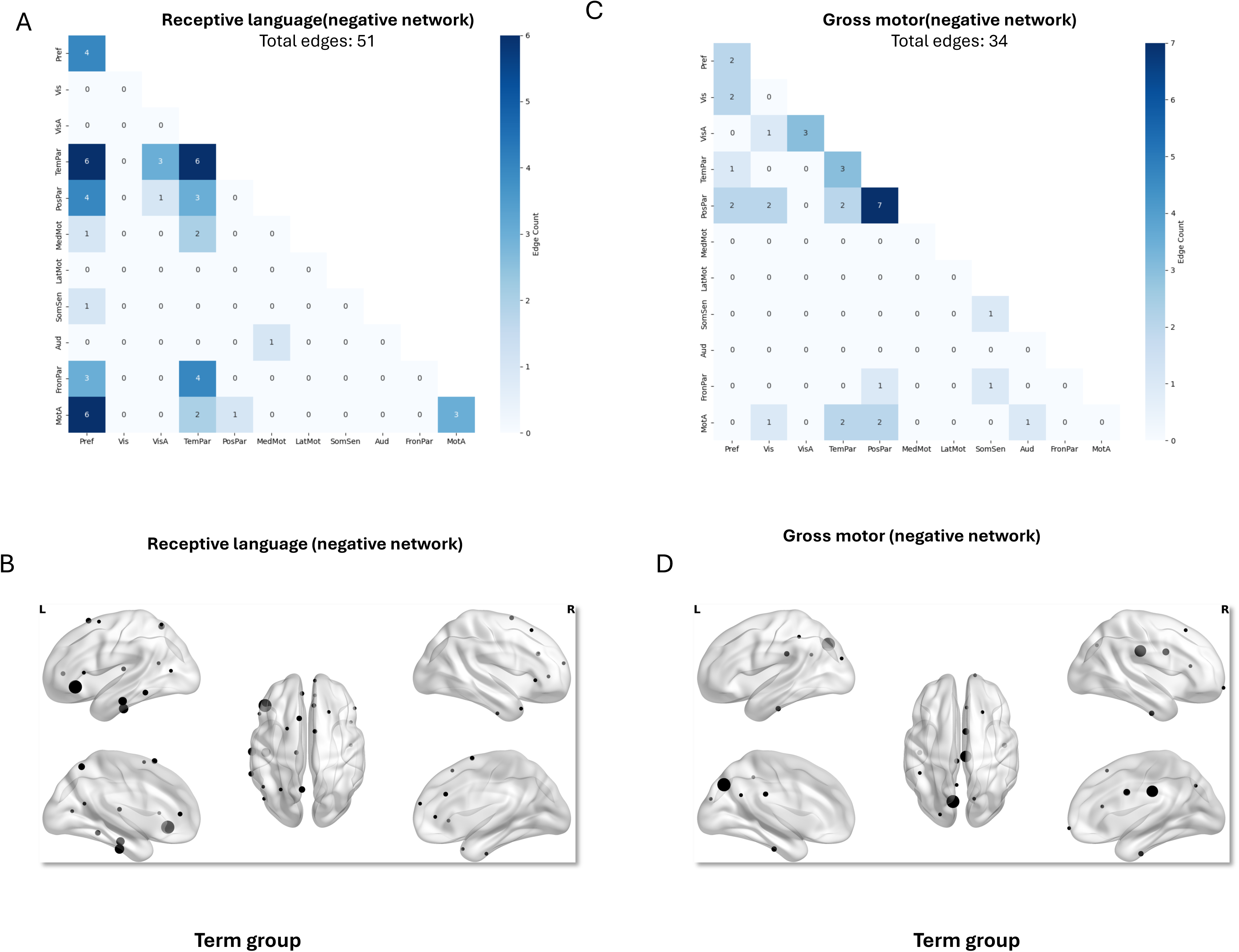
Negative networks: receptive language and gross motor predictive anatomy(term cohort). (A,C) Connections plotted as the number of edges within and between each pair of canonical networks for receptive language score and gross motor score respectively(left to right). (B,D) High-degree ROIs for the same models. Only ROIs with a degree ≥ one-sixth of the highest ROI are displayed; node size is proportional to degree.

### Cognition Prediction

The prediction model of cognition for the whole cohort was statistically significant (see Table 2); the model primarily included nodes located in the visual and auditory cortices, left precentral gyrus, right precuneus, right middle cingulate cortex, and right supplementary motor area (Figure 2B). The visual, auditory, and motor association networks played a predominant role in this prediction model, with the strongest connections observed between the visual network and the auditory, motor association, and medial motor networks(Figure 2A).

The model of cognition that included only term-born participants was also statistically significant (see Table 2). The nodes were primarily located in the visual cortex, left precentral gyrus, left precuneus, and right superior frontal gyrus, with the highest node degree observed in the visual cortex (Figure 2D). The visual, visual association, and motor association networks were highly involved in the prediction model, with the strongest connections observed between the visual network and the auditory, temporoparietal, and motor association networks. In addition, the motor association network demonstrated a high number of connections with the posterior parietal network (Figure 2C).

The cognition model that included only the pre-term born neonates was statistically significant (see Table 2). The cognitive–preterm prediction network exhibited the greatest concentration of nodes within the auditory cortex and the highest degree node was the left fusiform gyrus.

Additional nodes with lower connectivity were distributed across the right supplementary motor area, left parahippocampal gyrus, and left middle frontal gyrus (Figure 2F). The auditory and temporoparietal networks were highly involved, with the strongest connections observed within the auditory network and between the auditory and temporoparietal networks (Figure 2E).

### Language Prediction

The language prediction model with the whole cohort was statistically significant for the composite language score, expressive language score and receptive language score (see Table 2). Within visual and auditory cortex there were high degree nodes across all three language models (Figure 3A-3C), with the majority of edges dominated by the connection between visual network and auditory network in the three different language scores (Figure 4A-4C).

For the term-born group, the language prediction models were statistically significant for the composite language score, expressive language score, and receptive language score (see Table 2). Nodes in all three models were primarily located in the visual cortex, visual association cortex, and auditory cortex (Figure 3D-3F). The connectivity patterns across the three models were broadly similar, with the expressive language model exhibiting the greatest number of edges. Prominent connections in all three models included those between the visual network and the auditory and temporoparietal networks, between the visual association network and the auditory network, as well as within-network connections of the visual network (Figure 4D-4F). In addition, for the term group, a negative network associated with the receptive language score was identified (see Table 2). Nodes within this network were primarily located in the left inferior frontal gyrus, left inferior temporal gyrus, left middle temporal gyrus, and left precuneus (Figure 7B). The prefrontal and temporoparietal networks were core part of the prediction model, with prominent connections observed between the prefrontal network and the motor association and temporoparietal networks, as well as within the temporoparietal network itself (Figure 7A).

For the preterm-born group, the language prediction model was statistically significant for composite language score and receptive language score (see table 2). Nodes in both models were mainly located in the left fusiform gyrus, right lingual gyrus, and left superior temporal gyrus (Figure 3G-3H). The connectivity patterns for the composite language and receptive language models were similar to each other. In both the composite and receptive language models, the temporoparietal, auditory, and visual networks were actively involved, with strong connections observed between the temporoparietal network and auditory network (Figure 4G-4H).

### Motor prediction

Overall, motor prediction effects were weaker than those observed for cognitive and language outcomes. Permutation testing indicated only trend-level significance, and both mean validation accuracy and effect sizes (Cohen’s d_z_) were lower relative to the cognitive and language models (see table 2). In addition, the motor models involved substantially fewer predictive edges, suggesting a sparser and less robust underlying signal compared with cognition and language.

The motor prediction model with the whole cohort was statistically significant for composite motor score, fine motor score, and gross motor score (see table 2). Nodes in visual and auditory cortex were prominent in the composite and fine motor models, whereas nodes in auditory cortex were predominant in the gross motor model (Figure 5A-5C). Accordingly, connectivity between visual and auditory networks was prominent in the composite and fine motor models, while the connectivity between auditory and temporoparietal networks was predominant in gross motor model (Figure 6A-6C).

In the term-born group, the motor prediction model was statistically significant for composite motor score, and fine motor score (see table 2). Nodes were mainly in visual and auditory cortex in the composite motor model while in the fine motor model, the nodes were mainly in visual cortex, right anterior cingulate cortex, and right precentral gyrus (Figure 5D-5E). The connectivity between the visual and auditory networks played a dominant role in the composite motor network while the connectivity between the visual and medial motor networks was most prominent in the fine motor network (Figure 6D-6E). In addition, a negative model in the term cohort was statistically significant for gross motor score was statistically significant (see Table 2). Nodes were mainly located in the left precuneus and right middle cingulate cortex(Figure 7D). The posterior parietal network was highly involved, particularly with strong within-network connectivity(Figure 7C).

In the preterm-born group, the motor prediction models were statistically significant for the composite motor score and gross motor score (see Table 2). Nodes within the auditory cortex were prominent in both models, and the inferior temporal gyrus node showed a high degree of connectivity in both the composite and gross motor models. Compared with the composite motor model, the gross motor model included a greater number of nodes in the right precentral and postcentral gyri, left supramarginal gyrus, and right superior and inferior frontal gyri (Figure 5F-5G). The overall connectivity patterns were broadly similar in both models. The auditory network was highly active in both the composite and gross motor networks, showing strong within-network connections and robust links to the temporoparietal network. The motor association network was more prominently involved in the gross motor model than in the composite motor model (Figure 6F-6G).

### Model differences between term-born and preterm-born cohorts

To investigate the difference between term and preterm group, we first evaluated the relationship between average network strength and behaviour in the whole cohort, the term-born cohort, and the preterm-born cohort (Figure 8). It showed a steeper slope for the preterm cohort than for the term cohort (composite cognition: t(552)=-3.4, p=0.0008; composite language: t(552)=-4.8, p<0.0001; composite motor: t(552)=-1.2, p=0.22). Model anatomy also diverged by cohort: in term-born cohort, prediction relied most strongly on visual and auditory systems, whereas in preterm-born cohort temporoparietal and auditory systems were more prominently involved.

**Figure 8.**
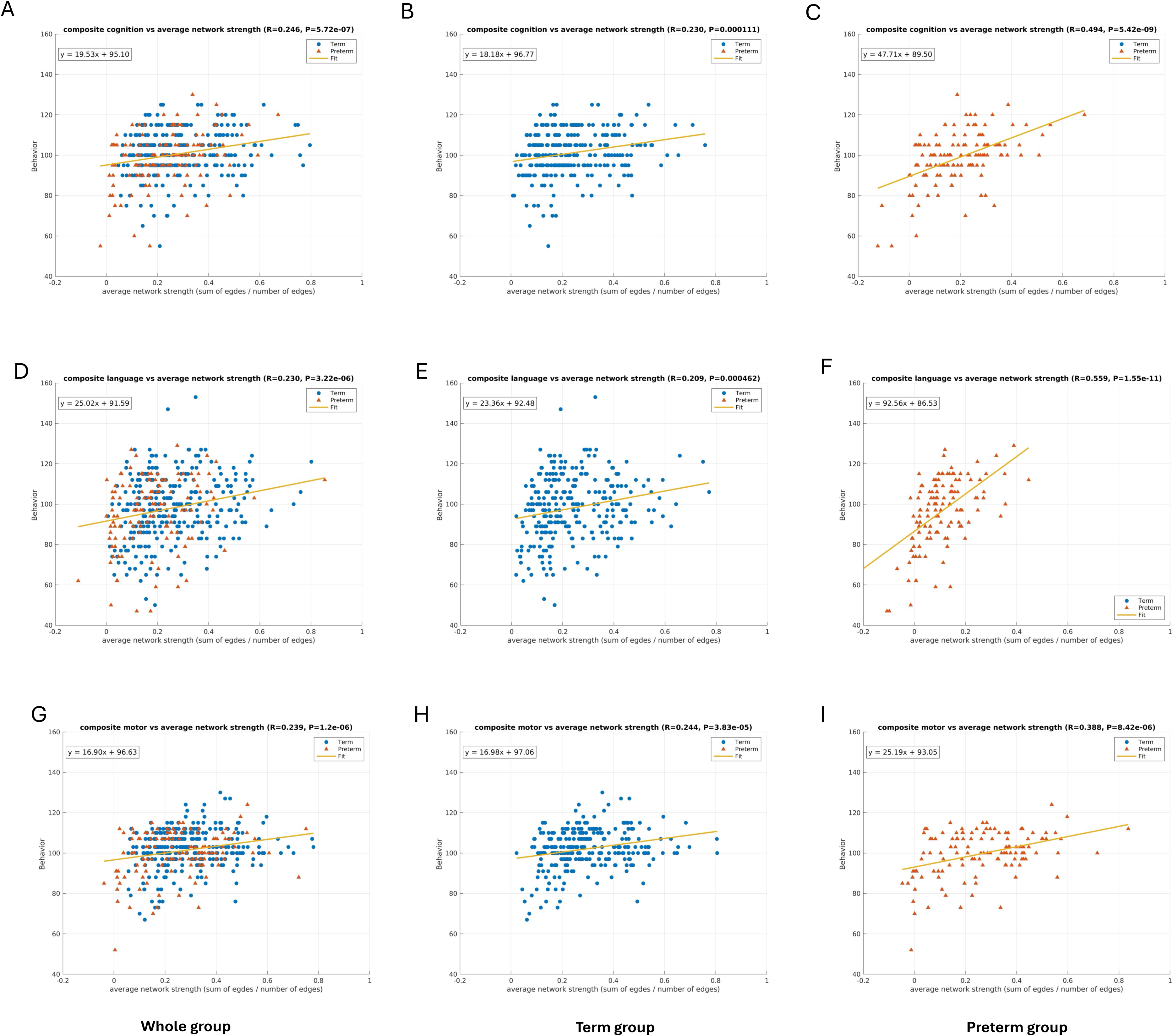
Average network strength versus behaviour by cohort. Scatterplots of average network strength(sum of edges/number of edges) versus Bayley-III composite scores are shown for whole-cohort, term-born cohort, and preterm-born cohort (left to right) across cognition, language, and motor composite scores (top to bottom).

As the next step, we then examined for statistical differences between interhemispheric and intrahemispheric connectivity in term and preterm cohort. In the preterm group, a one-sided binomial test showed that the number of interhemispheric edges exceeded that of intrahemispheric edges for every Bayley-III measure except gross motor (p < 0.001), while the term group showed similar inter- and intra-hemispheric edge counts (Figure 9D). The network sources of this hemispheric pattern were consistent across outcomes. In cognition and language models of preterm cohort, interhemispheric connectivity was primarily contributed by auditory and temporoparietal networks, whereas in the motor-preterm model the auditory network showed the strongest interhemispheric emphasis(Figure 9A-9C); this pattern persisted for gross motor(see S17D). Figure 9 summarizes cognition, language, and motor composite results; results for the language subscales (receptive, expressive) and the motor subscales (gross, fine) are provided in S17.

**Figure 9.**
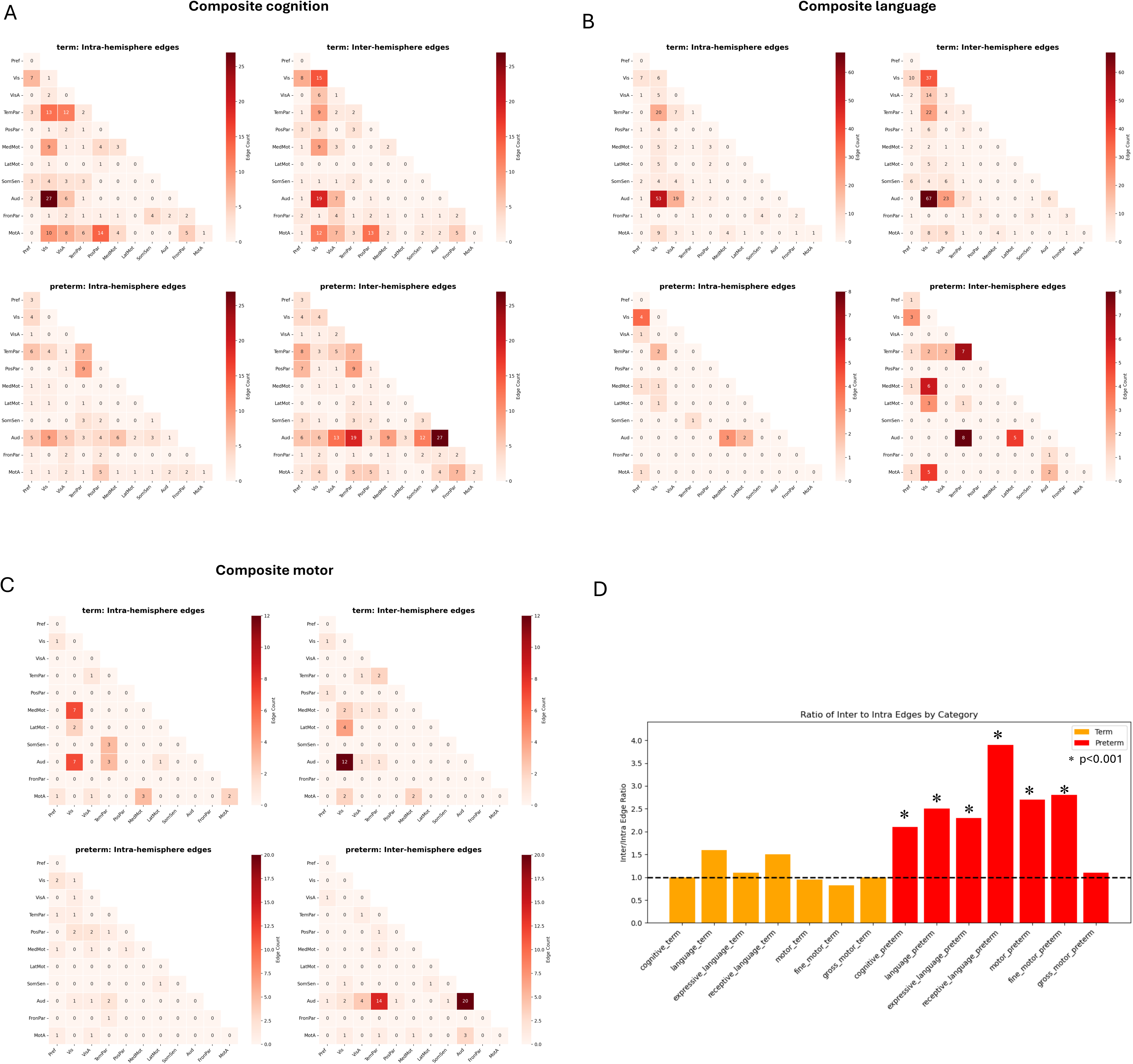
Inter-/intra-hemispheric connectivity patterns in term and preterm cohorts. (A,B,C) inter and intra-hemisphere between/within-network connectivity profiles of term-born and preterm-born cohorts for cognition, language and motor composite scores respectively; (D) Ratios of inter-hemispheric edges counts to intra-hemispheric edge counts of term-born and preterm-born cohorts for cognition, language, and motor models (including motor and language subscales). The asterisk (*) indicates that the number of inter-hemispheric edges is significantly greater than the number of intra-hemispheric edges.

### Integration of Term- and Preterm-Typical Connectivity Patterns in Whole-Cohort Models

Having established a cohort difference, we next found that in the whole cohort, predictive models reflected contributions from both term- and preterm-associated connectivity patterns. For example examining the cognition model, in the term group, predictive ROIs were concentrated in the left and right visual cortices, with additional contribution from the left precentral gyrus. In the preterm group, predictive ROIs were concentrated in the left auditory cortex, the left fusiform gyrus, and the right supplementary motor area. The whole-cohort cognition model contained ROIs from both term- and preterm-typical sets, reflecting both visual/visual-association components from term group and auditory components from preterm group within a single network. The same pattern was also evident for motor. The whole-cohort ROI set combined right visual cortex nodes typical of the term model with left auditory cortex nodes typical of the preterm model. And the connectivity pattern of the whole-cohort model integrated contributions from both cohorts—visual network emphasized in the term group and auditory network emphasized in the preterm group. For language, the whole-cohort network was similar to the term pattern but still included nodes characteristic of the preterm group, specifically nodes in right lingual gyrus and left superior temporal gyrus of the preterm group. Consistent with this interpretation, when we computed average network strength in the whole cohort using edges derived separately from the term-only and preterm-only models, both sets of edges showed significant correlations with behaviour (p < 0.001) (Figure S17). This indicates that the whole-cohort model contains contributions from both term-typical and preterm-typical connectivity patterns.

## Discussion

This study demonstrated that neonatal resting-state functional connectivity, when analysing through a stability-driven, ROI-constrained connectome-based predictive modeling (CPM) framework, can predict cognitive, language, and motor outcomes at 18 months of age. We identify distinct yet complementary network signatures between term and preterm neonates: models based on term cohort relied primarily on visual-auditory integration, whereas models based on preterm cohort depended more on auditory-temporoparietal connectivity with pronounced interhemispheric bias. Whole-cohort models reflected combined connectivity signatures of term and preterm infants. These findings refine our understanding of early connectomic organization and its link to neurodevelopmental trajectories.

### Early sensory foundations of behavioural prediction

Our results demonstrated that neonatal functional connectivity within and between primary sensory cortices—particularly visual and auditory systems—robustly predicted later cognitive, language, and motor performance at 18 months. This was consistent with results that these systems exhibit coherent organization at and before term age (Doria et al., 2010; Fransson et al., 2009), but extended it by demonstrating predictive power for individual neurodevelopmental outcomes. These findings reinforced the concept of a sensory-first developmental hierarchy, where early-maturing sensorimotor systems provide the neural scaffolding upon which higher-order cognition later emerges (Gao et al., 2015).

### Divergent developmental pathways in term and preterm neonates

A major contribution of this study lies in identifying distinct predictive architectures between term and preterm cohorts. In term-born neonates, predictive edges prominently link visual and auditory cortices, reflecting emerging multisensory integration. In contrast, preterm-born neonates rely primarily on auditory and temporoparietal systems, with pronounced interhemispheric connectivity. Compared to term infants, visual-auditory interaction doesn’t seem to contribute to the predictive model in preterm infants. This divergence likely mirrors asynchronous maturation: the auditory system matures earlier and is relatively resilient to prematurity (Angrisani et al., 2014), whereas the visual system—especially dorsal visual and visuomotor pathways—undergoes accelerated but vulnerable postnatal development (Atkinson & Braddick, 2011; Braddick et al., 2011; Loenneker et al., 2011). The absence of audiovisual coupling in preterm models suggests a disrupted multisensory convergence window, consistent with behavioral reports of reduced audiovisual speech perception(Berdasco-Muñoz et al., 2019; Imafuku et al., 2019) and impaired multisensory processes persisting into school age(Décaillet et al., 2024) in preterm .

The increased interhemispheric connectivity observed in preterm networks (Naoi et al., 2013; Wilke et al., 2014) aligns with prior reports of altered bilateral coordination in preterm infants, although its functional significance—whether compensatory or otherwise—remains unclear.

Rather than forming efficient intrahemispheric circuits, preterm neonates may rely on diffuse bilateral synchronization to sustain global integration—an organization that may reflect delayed segregation and reduced efficiency typical of early network immaturity (Bouyssi-Kobar et al., 2019; López-Guerrero & Alcauter, 2025). Such connectivity may preserve short-term function but predict later vulnerability in executive or visuospatial domains.

### Domain-specific insights Cognition

Cognitive predictions drew primarily on visual, auditory, and medial motor cortices, with minimal prefrontal involvement. The negligible contribution of prefrontal regions to prediction likely reflects their structural and functional immaturity in the neonatal period (Tsujimoto, 2008). Prefrontal–parietal and prefrontal–temporal connectivity strengthen rapidly after the first months of life through experience-dependent processes, including caregiver interaction and environmental enrichment (Hodel, 2018; Sheridan & McLaughlin, 2014). Thus, early predictive signals appear dominated by systems whose organization is innate or sensorily driven, whereas prefrontal contributions may emerge only after sufficient experiential scaffolding.

### Language

In language prediction, visual network was strongly engaged in term infants and remained involved even in preterm infants—unlike cognition and motor models, where visual networks were absent. This highlights the unique importance of visual input in early language development. Infants integrate visual cues such as lip movements, facial expressions, and gaze with auditory information to support speech perception and word learning (K et al., 2015; Michon et al., 2022; Putzar et al., 2010). These findings suggest that the visual system provides a fundamental and resilient scaffold for early language acquisition.

### Motor

For motor outcomes, fine motor prediction in term cohort involved strong visual–medial motor connectivity, while gross motor prediction relied on medial motor and temporoparietal networks. This dissociation may reflect domain-specific visuomotor and postural systems: fine motor skills depend on early maturation of occipital–motor circuits supporting hand–eye coordination, whereas gross motor control engages body-centered parietal–motor integration critical for whole-body movement and balance.

### Methodological and translational implications

Methodologically, this work advances connectome-based predictive modeling by introducing a stability- and hub-oriented framework that enhances both robustness and interpretability. We demonstrate that predictive accuracy is highest for connections linked to high-degree ROIs and progressively declines as lower-degree regions are added—a pattern explained by a corresponding decrease in signal-to-noise ratio. Permutation testing confirmed that the observed predictive performance was statistically significant, underscoring the reliability of the extracted signal. This hub-constrained CPM thus reveals connectivity–behavior relationships with small effects that are often obscured in conventional CPM frameworks. And from independent validation we demonstrate signal edges from lower-degree regions also carry predictive information. Beyond neonatal imaging, the same principle—anchoring prediction to stable, high-degree network nodes—may generalize to other developmental or clinical fMRI datasets where low SNR and participant variability often limit predictive reliability. Future work could extend this approach by incorporating complementary graph metrics (e.g., betweenness or eigenvector centrality) or dynamic connectivity measures to capture temporal variations in early brain network organization.

Clinically, the identification of distinct sensory predictors in preterm infants supports the design of targeted early interventions. Programs emphasizing enriched visual–auditory co-stimulation (Tang et al., 2025) could promote more balanced sensory integration, potentially mitigating downstream cognitive and language delays. The feasibility of using rs-fMRI to stratify risk at birth offers an avenue for precision neurodevelopmental monitoring.

### Limitations and future directions

Despite robust validation, several limitations warrant consideration. First, subcortical signal attenuation limits our inference about basal ganglia and thalamic contributions, despite their critical role in sensorimotor integration. Second, the restricted variance of Bayley-III scores in this relatively healthy cohort may underestimate predictive sensitivity and reduce sensitivity to detect finer-grained functional differences.

Future research could expand to clinical populations and longitudinally integrate genetic and environmental factors (e.g., neonatal nutrition, parental interaction) to clarify the causal pathways linking early brain connectivity to later cognitive outcomes. Further investigations could incorporate dynamic network analyses and multimodal imaging (e.g., structural MRI, diffusion imaging, EEG) to elucidate how functional and structural coupling evolves with sensory experience. Combining connectome-based predictive modeling (CPM) with graph-theoretic efficiency metrics or generative modeling could provide deeper insights into the mechanistic principles underlying adaptive and maladaptive developmental trajectories.

## Conclusion

This study identifies neonatal sensory networks as foundational predictors of cognitive, language, and motor development and uncovers distinct term–preterm connectomic architectures. Visual–auditory integration emerges as a hallmark of typical maturation, whereas preterm infants may exhibit compensatory reliance on auditory and interhemispheric pathways. Together, these findings provide mechanistic insight into the neural basis of early developmental risk and lay the groundwork for neuroimaging-informed early intervention strategies.

## Acknowledgement

All calculations were performed on the high performance cluster maintained by the Trinity College Research IT unit. The research was supported by funding through a joint scholarship programme of the China Scholarship Council and Trinity College Dublin. Data used in the preparation of this manuscript were obtained from the National Institute of Mental Health (NIMH) Data Archive (NDA). NDA is a collaborative informatics system created by the National Institutes of Health to provide a national resource to support and accelerate research in mental health. Dataset identifier(s): 10.15154/vwwg-kc17. This manuscript reflects the views of the authors and may not reflect the opinions or views of the NIH or of the Submitters submitting original data to NDA.

## Author Contributions

Arun L. W. Bokde conceptualized the application of CPM and supervised the project, and Mi Zou developed the novel R_CPM methodology, performed the data analysis, and wrote and revised the manuscript. Arun L. W. Bokde contributed to language polishing.

