## Supplemental Data 1 for "Neonatal sensory networks at birth predict cognitive, language, and motor outcomes at 18 months"

Table 1 Correlation between prediction accuracy and SNR

| | Whole cohort | Term cohort | Preterm cohort |
|---|---|---|---|
| Composite cognition | 0.96 | 0.87 | 0.7 |
| Composite language | 0.8 | 0.84 | 0.62 |
| Expressive language | 0.88 | 0.95 | 0.65 |
| Receptive language | 0.85 | 0.85 | 0.6 |
| Composite motor | 0.76 | 0.84 | 0.55 |
| Fine motor | 0.4 | 0.91 | 0.39 |
| Gross motor | 0.56 | 0.88 | 0.87 |

Table 2 Coverage with highset validation accuracy in preterm cohort

| preterm cohort | best coverage | T(299) | p |
|---|---|---|---|
| composite cognition | 90% | 2.05 | 0.041004915 |
| receptive language | 50% | 4.08 | 5.77E-05 |
| composite motor | 30% | 8.12 | 1.21E-14 |

Note: The best coverage refers to the coverage level that achieves the highest validation accuracy compared to the full coverage (100%). *T(299)* represents the paired *t*-test statistic comparing the best coverage and the 100% coverage across 300 data splits. For all behaviors not shown here, the best coverage corresponds to 100% (all statistics comparing coverage 100% and every intermediate coverage—10%-90%—see coverage_t_test.xlsx).

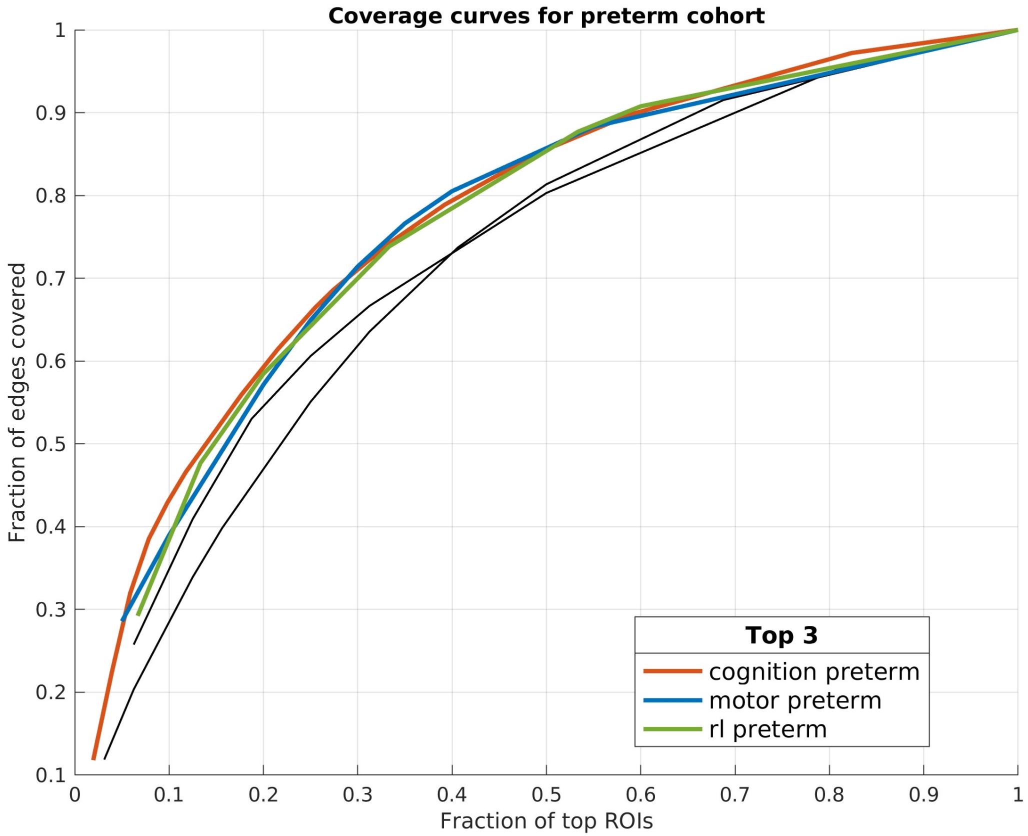


**Figure S1. Coverage curves for the preterm cohort.**
Coverage curves show the relationship between the fraction of top-ranked ROIs (number of top ROIs/number of total ROIs) and the fraction of total network edges they cover (number of edges/total edges). Each colored line represents one of the top three behavioral domains—composite cognition (orange), composite motor (blue), and receptive language(rl) (green)—for the preterm cohort. The colored curves indicate that a small subset of top ROIs captures a large proportion of network edges. The thinner black curves correspond to other behavioral models not in the top three. This corresponds to the three behaviors in table 2 showed best coverage lower than 100%. In summary, except for cases in which a small fraction of top ROIs accounts for a large fraction of the total edges, the overall SNR increases as the signal edges of low degree ROIs were added.

### Composite cognition (preterm)

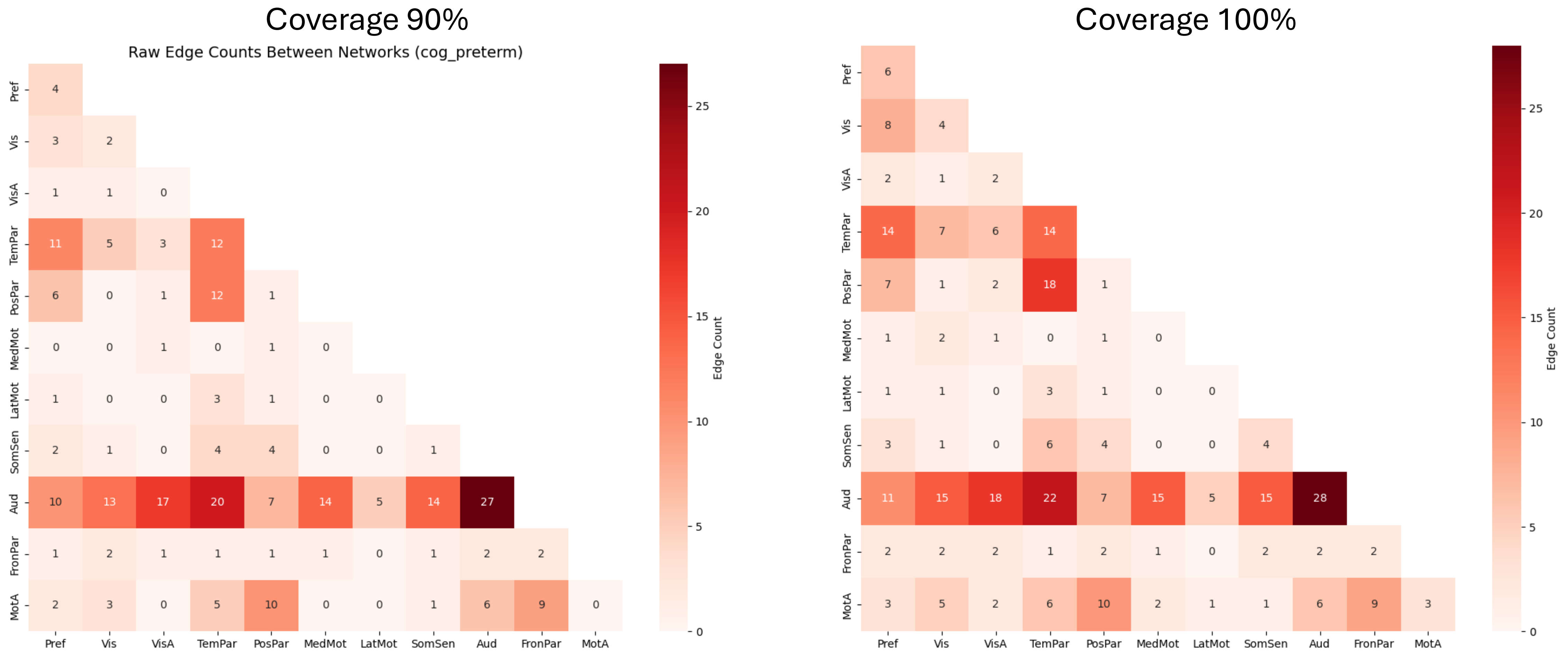


**Figure S2. Comparison between best coverage and full coverage (100%).**
The figure compares best coverage (the coverage level that achieved the highest validation accuracy) with the full coverage condition (100%). The connectivity pattern stays consistent in the two thresholds.

### Receptive language (preterm)

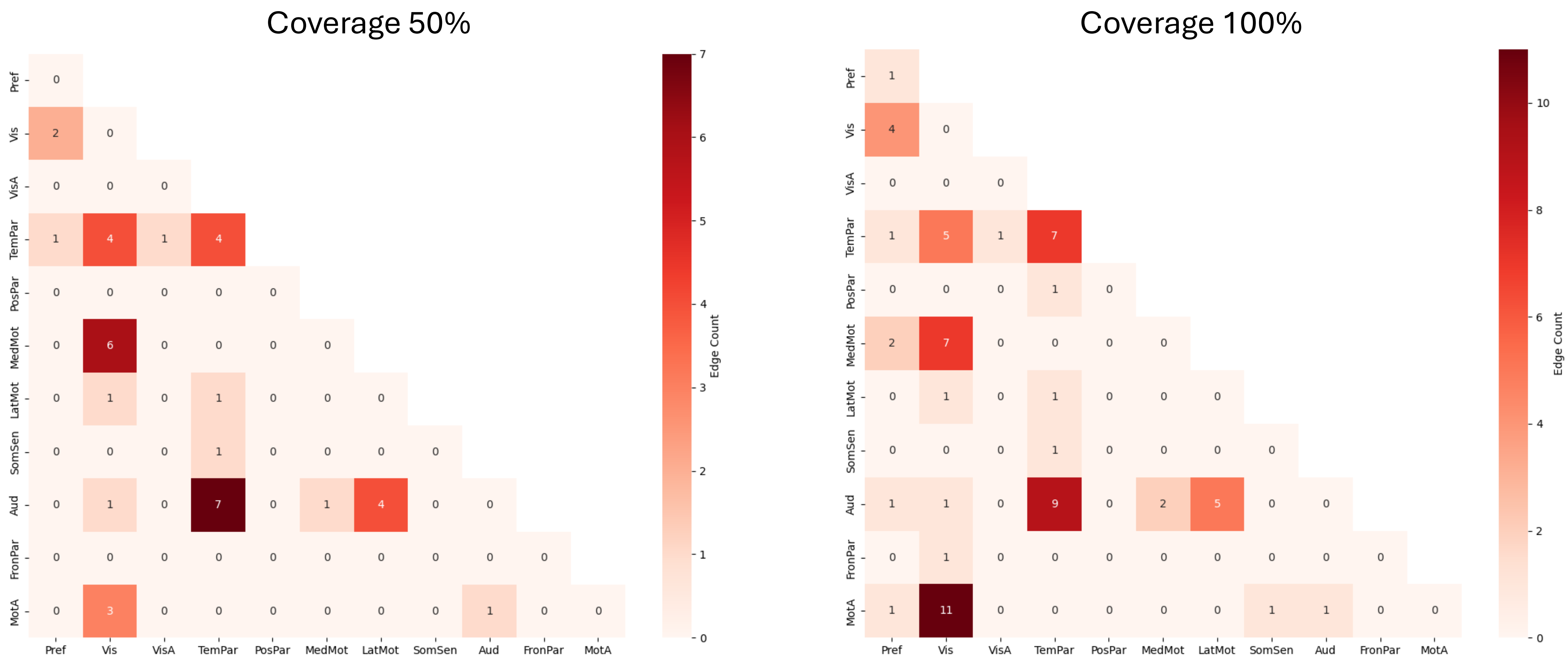


**Figure S3. Comparison between best coverage and full coverage (100%).**
The figure compares best coverage (the coverage level that achieved the highest validation accuracy) with the full coverage condition (100%). The connectivity pattern stays consistent in the two thresholds.

### Composite motor (preterm)

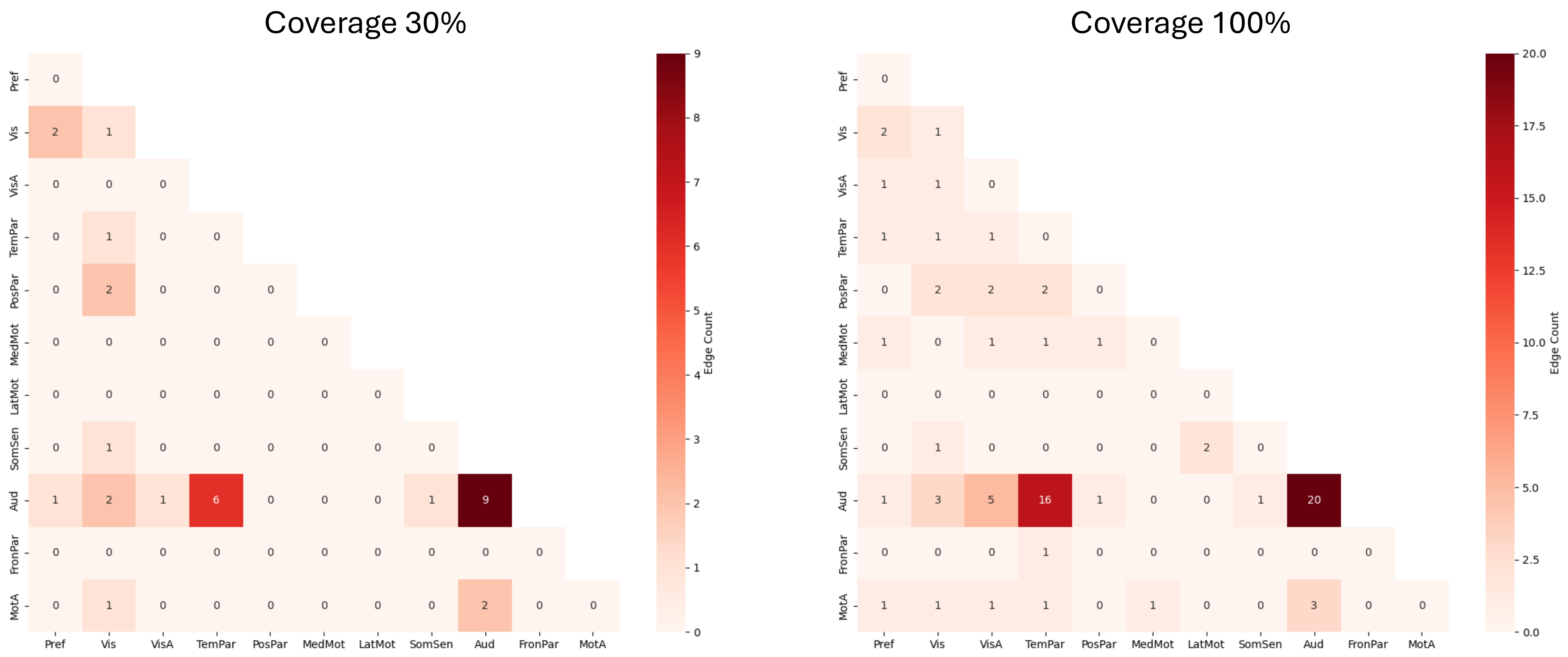


**Figure S4. Comparison between best coverage and full coverage (100%).**
The figure compares best coverage (the coverage level that achieved the highest validation accuracy) with the full coverage condition (100%). The connectivity pattern stays consistent in the two thresholds.

Table 3 Coverage with highset validation accuracy in term cohort

| term cohort | best coverage | T | p |
|---|---|---|---|
| composite language | 0.70 | 7.04 | 1.32E-11 |
| expressive language | 0.60 | 12.6<br>4 | 1.20E-29 |
| receptive language | 0.70 | 4.81 | 2.38E-06 |

Note: The best coverage refers to the coverage level that achieves the highest validation accuracy compared to the full coverage (100%). *T(299)* represents the paired *t*-test statistic comparing the best coverage and the 100% coverage across 300 data splits. For all behaviors not shown here, the best coverage corresponds to 100% (all statistics comparing coverage 100% and every intermediate coverage—10%-90%—see coverage_t_test.xlsx).

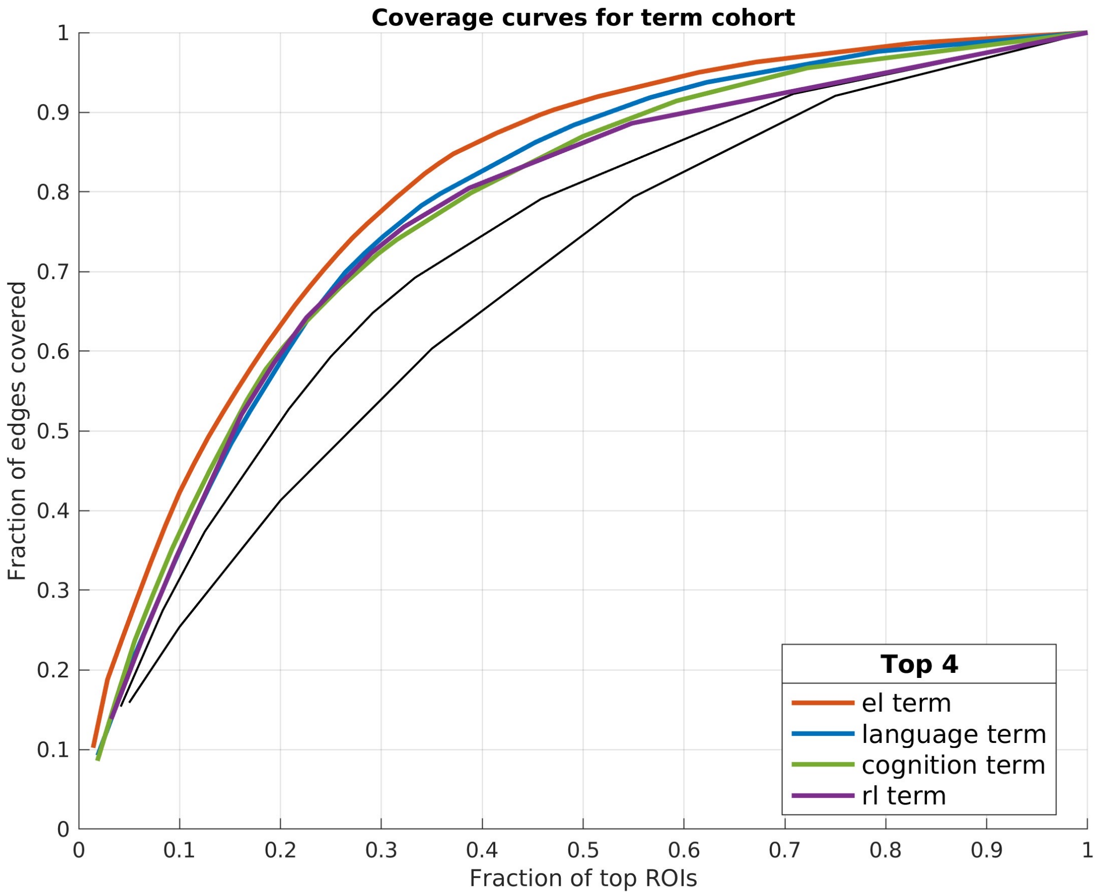


**Figure S5. Coverage curves for the term cohort.**
Coverage curves show the relationship between the fraction of top-ranked ROIs (number of top ROIs/number of total ROIs) and the fraction of total network edges they cover (number of edges/total edges). Each colored line represents one of the top four behavioral domains—expressive language(orange), composite language(blue), composite cognition(green), and receptive language(purple)—for the term cohort. The colored curves indicate that a small subset of top ROIs captures a large proportion of network edges. The thinner black curves correspond to other behavioral models not in the top three. This corresponds to the three behaviors in table 2 showed best coverage lower than 100%. In summary, except for cases in which a small fraction of top ROIs accounts for a large fraction of the total edges, the overall SNR increases as the signal edges of low degree ROIs were added.

### Composite language (term)

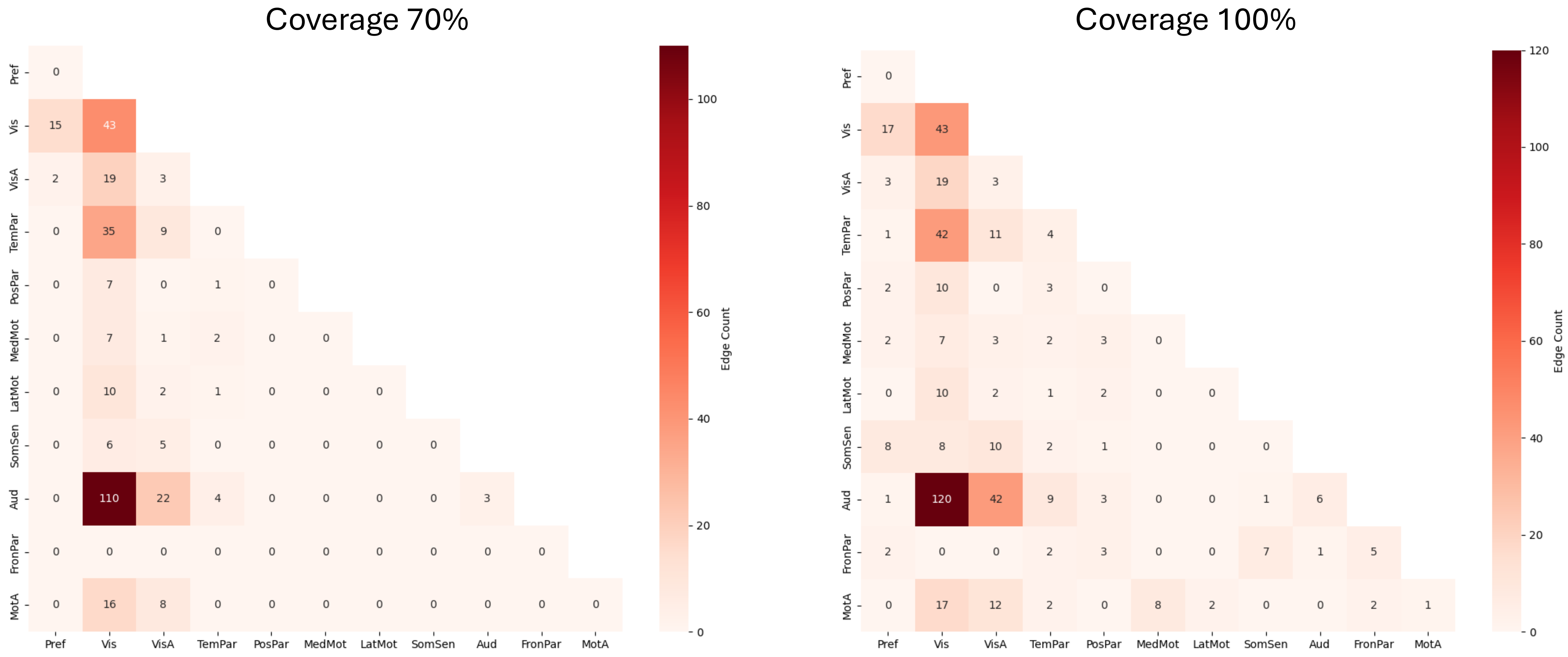


**Figure S6. Comparison between best coverage and full coverage (100%).**
The figure compares best coverage (the coverage level that achieved the highest validation accuracy) with the full coverage condition (100%). The connectivity pattern stays consistent in the two thresholds.

#### Expressive language (term)

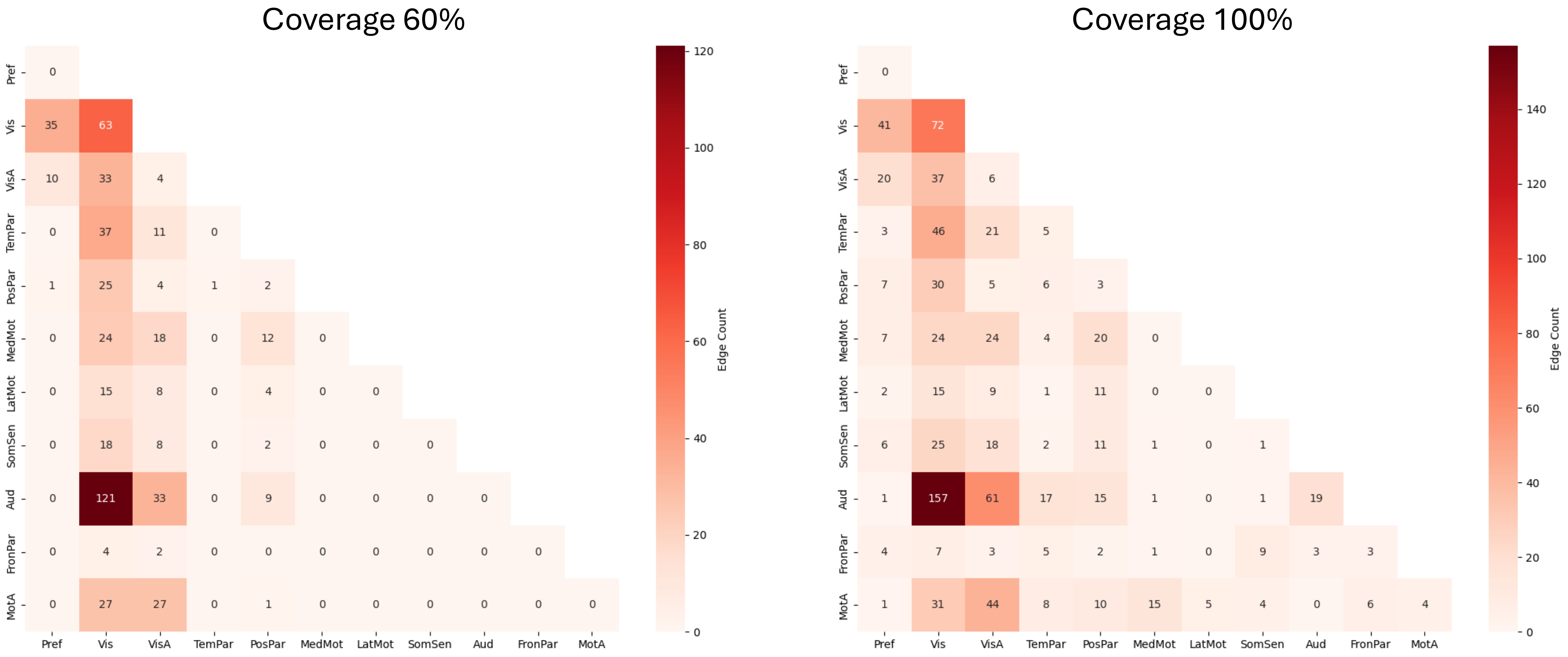


**Figure S7. Comparison between best coverage and full coverage (100%).**
The figure compares best coverage (the coverage level that achieved the highest validation accuracy) with the full coverage condition (100%). The connectivity pattern stays consistent in the two thresholds.

### Receptive language (term)

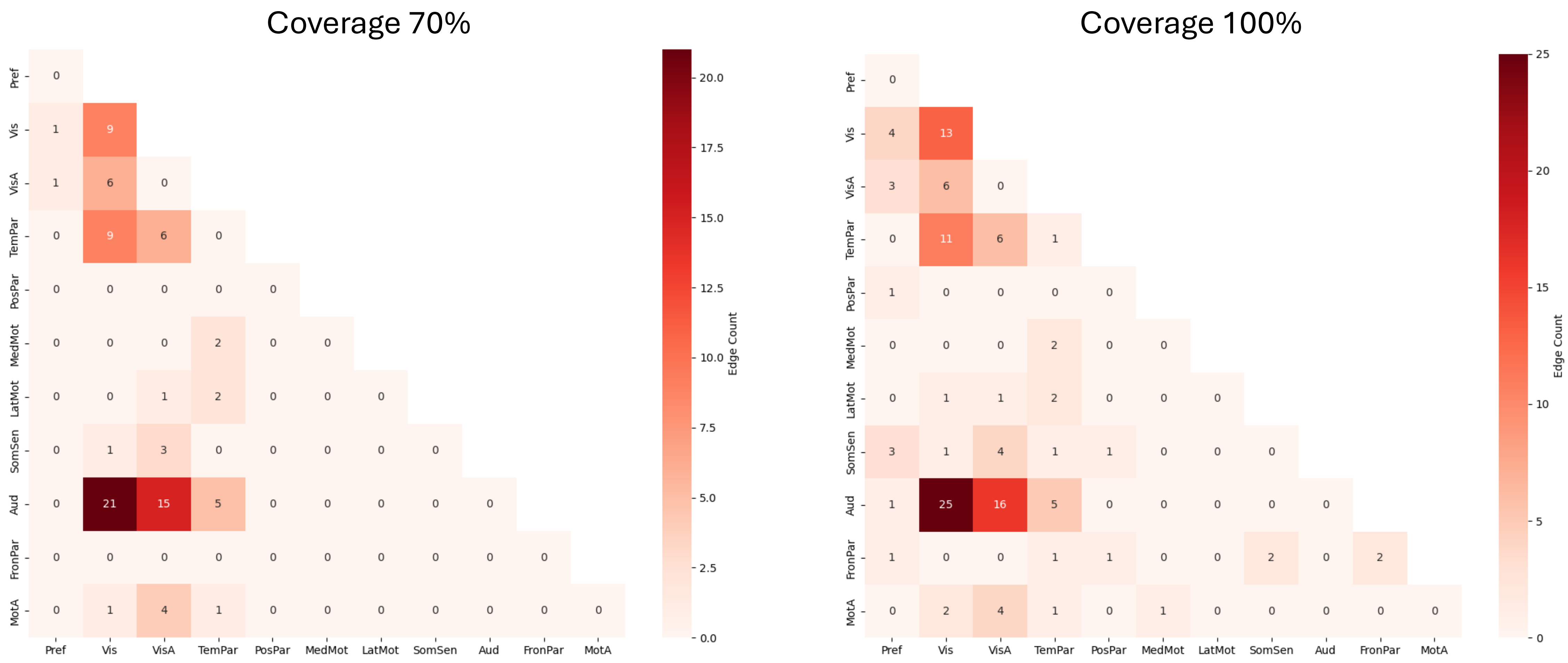


**Figure S8. Comparison between best coverage and full coverage (100%).**
The figure compares best coverage (the coverage level that achieved the highest validation accuracy) with the full coverage condition (100%). The connectivity pattern stays consistent in the two thresholds.

Table 4 Coverage with highset validation accuracy in whole cohort

| whole cohort | best coverage | T | p |
|---|---|---|---|
| composite language | 0.90 | 5.16 | 4.59E-07 |
| expressive language | 0.80 | 9.74 | 1.21E-19 |
| receptive language | 0.80 | 2.76 | 0.006049514 |
| composite motor | 0.80 | 3.17 | 0.001655323 |
| fine motor | 0.70 | 6.18 | 2.14E-09 |
| gross motor | 0.80 | 3.54 | 0.000469496 |

Note: The best coverage refers to the coverage level that achieves the highest validation accuracy compared to the full coverage (100%). *T(299)* represents the paired *t*-test statistic comparing the best coverage and the 100% coverage across 300 data splits. For all behaviors not shown here, the best coverage corresponds to 100% (all statistics comparing coverage 100% and every intermediate coverage—10%-90%—see coverage_t_test.xlsx).

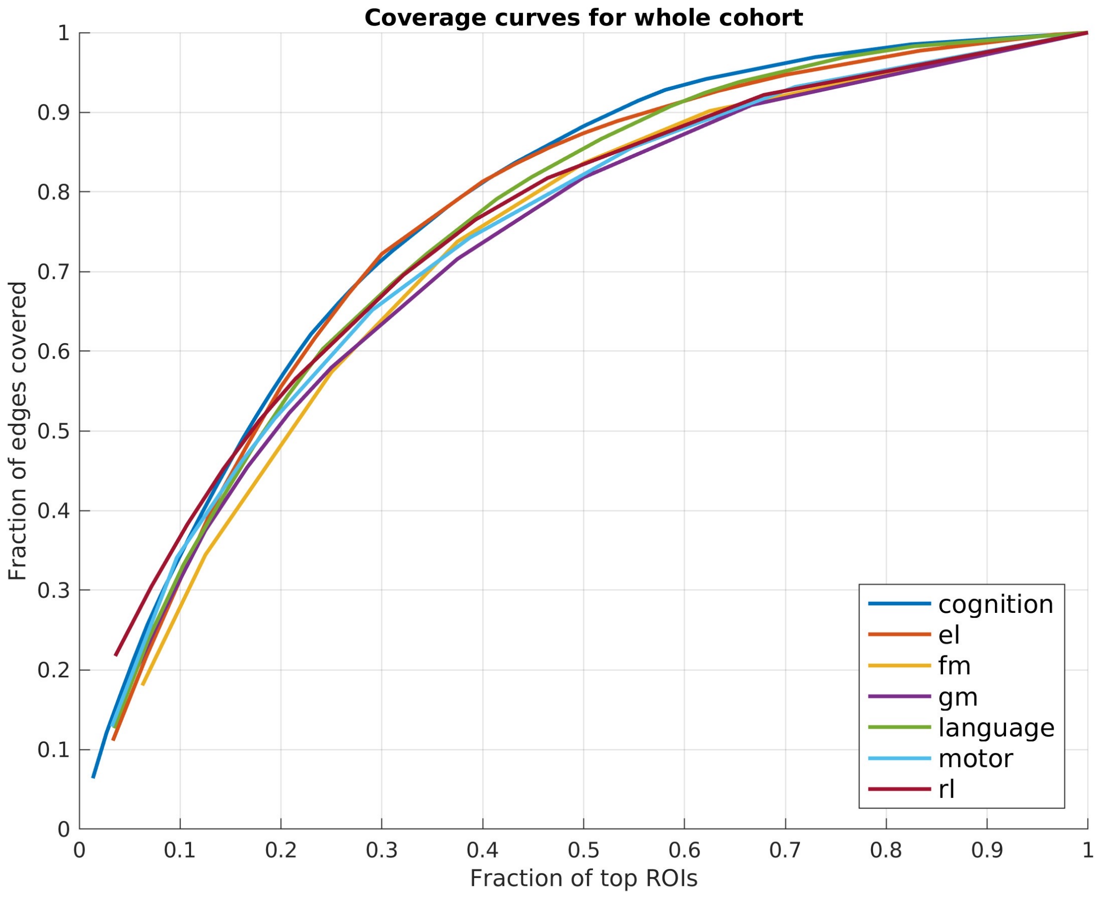


**Figure S9. Coverage curves for the whole cohort.**
Coverage curves show the relationship between the fraction of top-ranked ROIs (number of top ROIs/number of total ROIs) and the fraction of total network edges they cover (number of edges/total edges). There are no obvious visual difference between these lines, which corresponds to table 2, best coverage of all behaviors are close to each other.

### Composite language (whole)

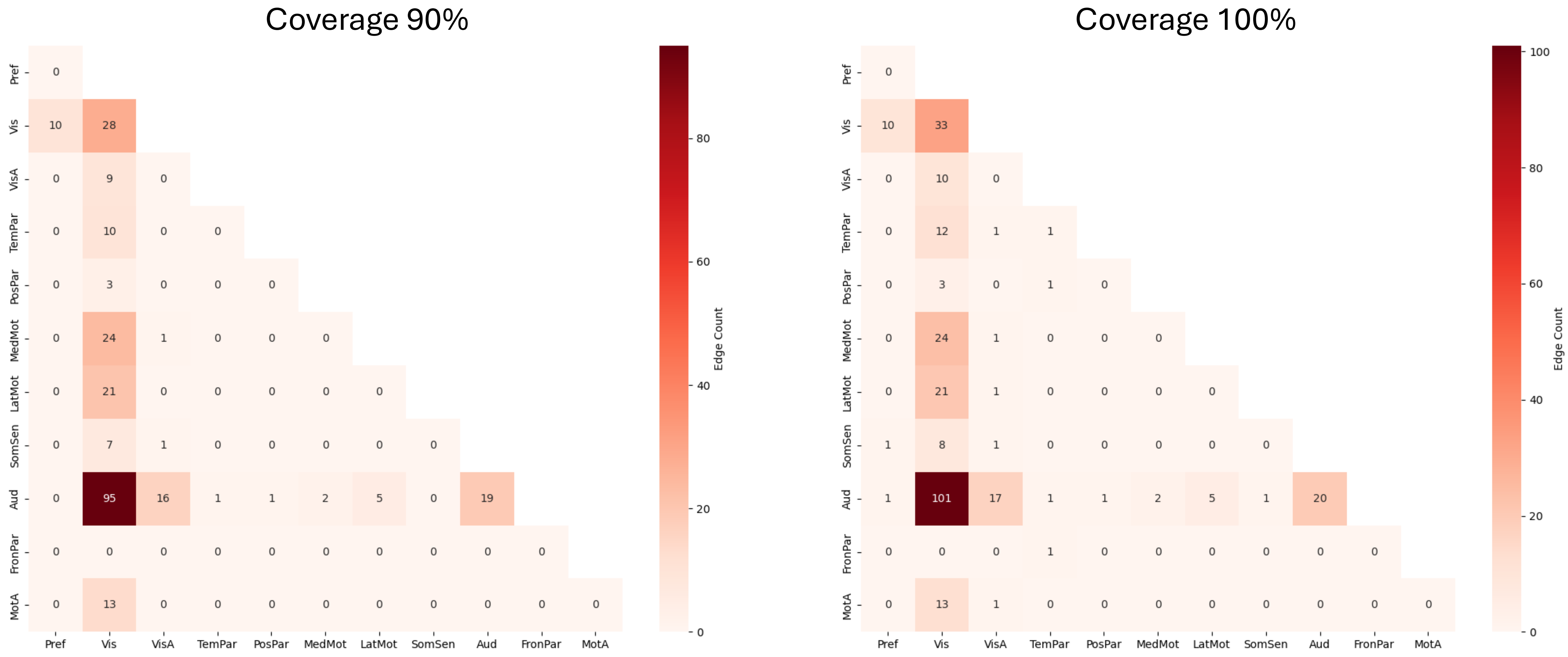


**Figure S10. Comparison between best coverage and full coverage (100%).**
The figure compares best coverage (the coverage level that achieved the highest validation accuracy) with the full coverage condition (100%). The connectivity pattern stays consistent in the two thresholds.

#### Expressive language (whole)

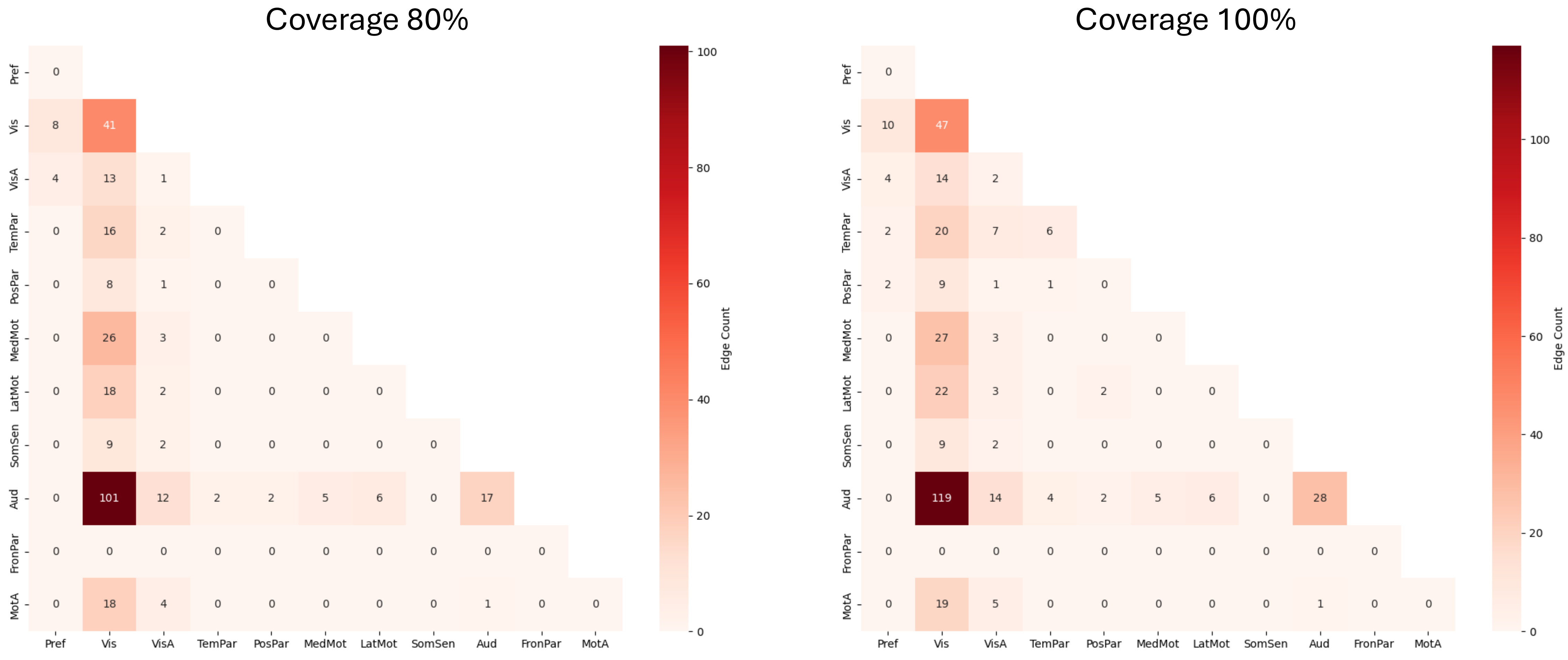


**Figure S11. Comparison between best coverage and full coverage (100%).**
The figure compares best coverage (the coverage level that achieved the highest validation accuracy) with the full coverage condition (100%). The connectivity pattern stays consistent in the two thresholds.

### Receptive language (whole)

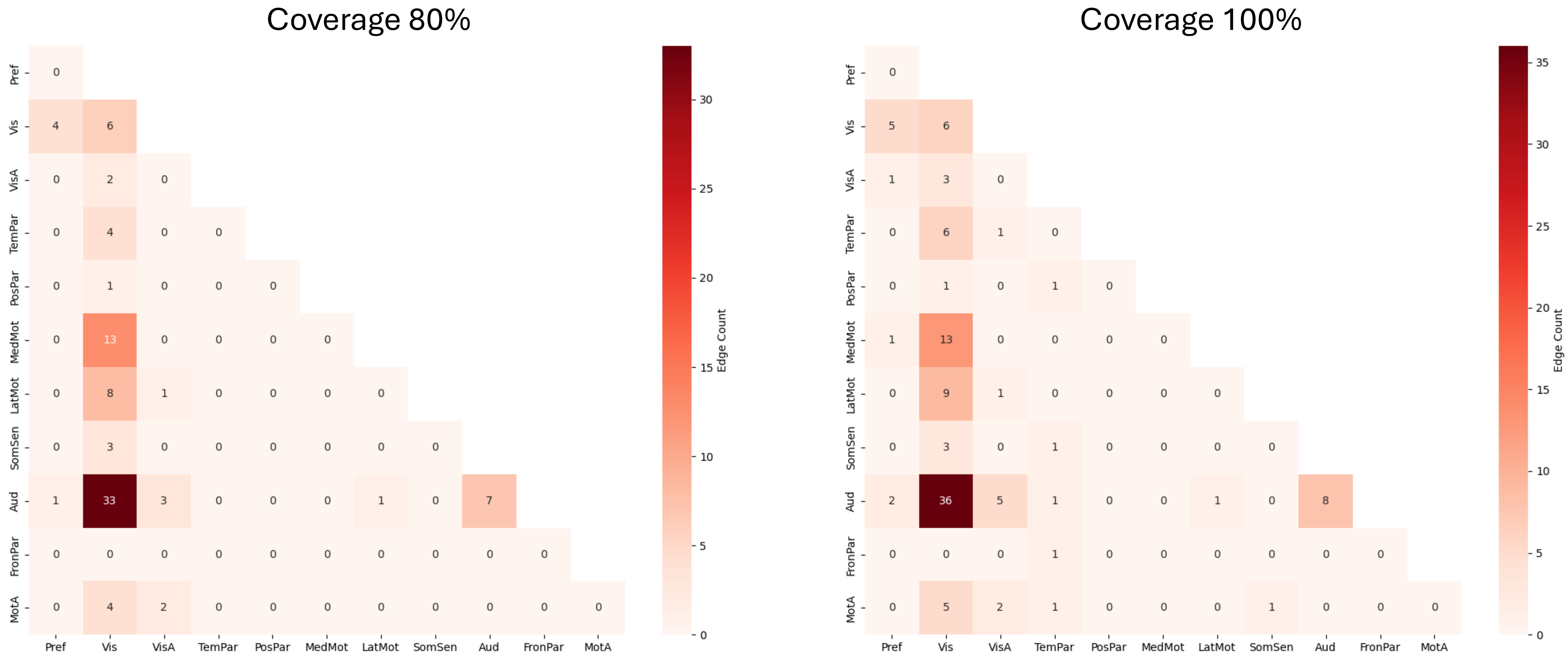


**Figure S12. Comparison between best coverage and full coverage (100%).**
The figure compares best coverage (the coverage level that achieved the highest validation accuracy) with the full coverage condition (100%). The connectivity pattern stays consistent in the two thresholds.

### Composite motor (whole)

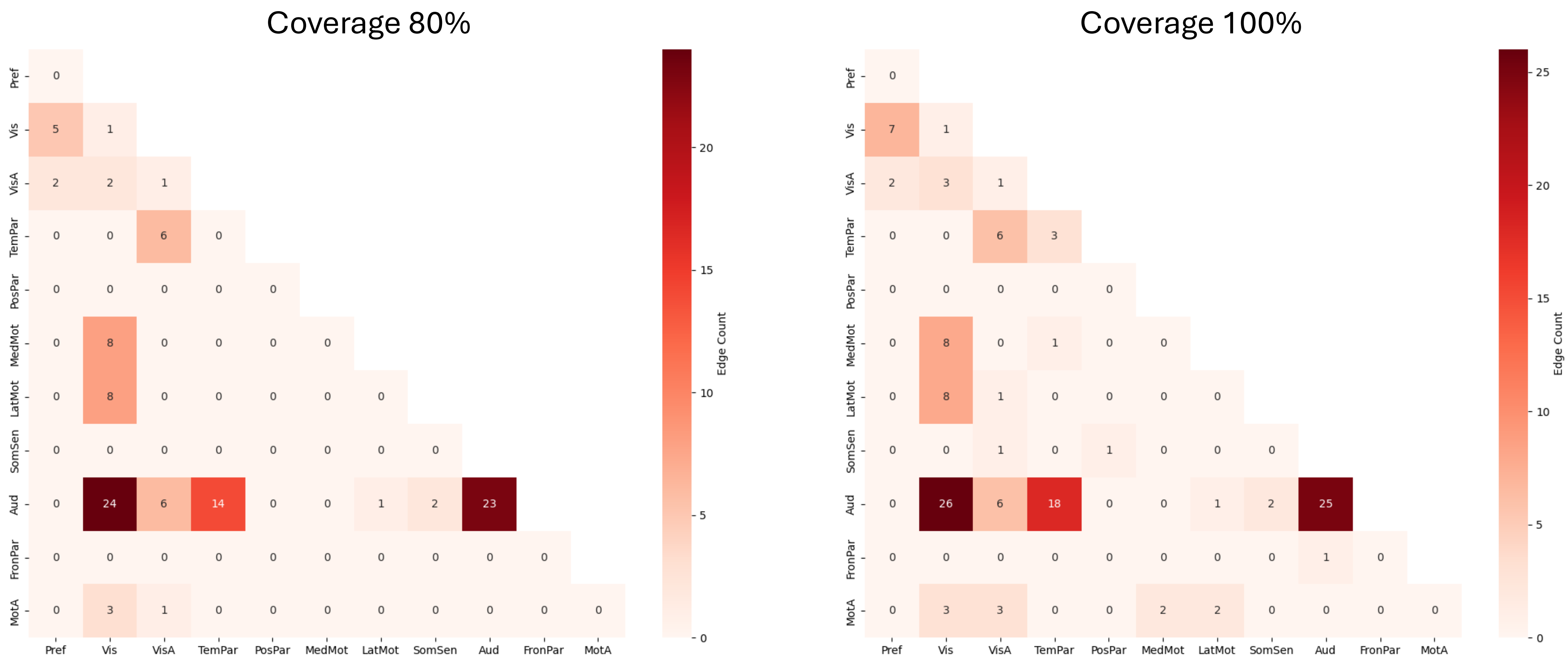


**Figure S13. Comparison between best coverage and full coverage (100%).**
The figure compares best coverage (the coverage level that achieved the highest validation accuracy) with the full coverage condition (100%). The connectivity pattern stays consistent in the two thresholds.

#### Fine motor (whole)

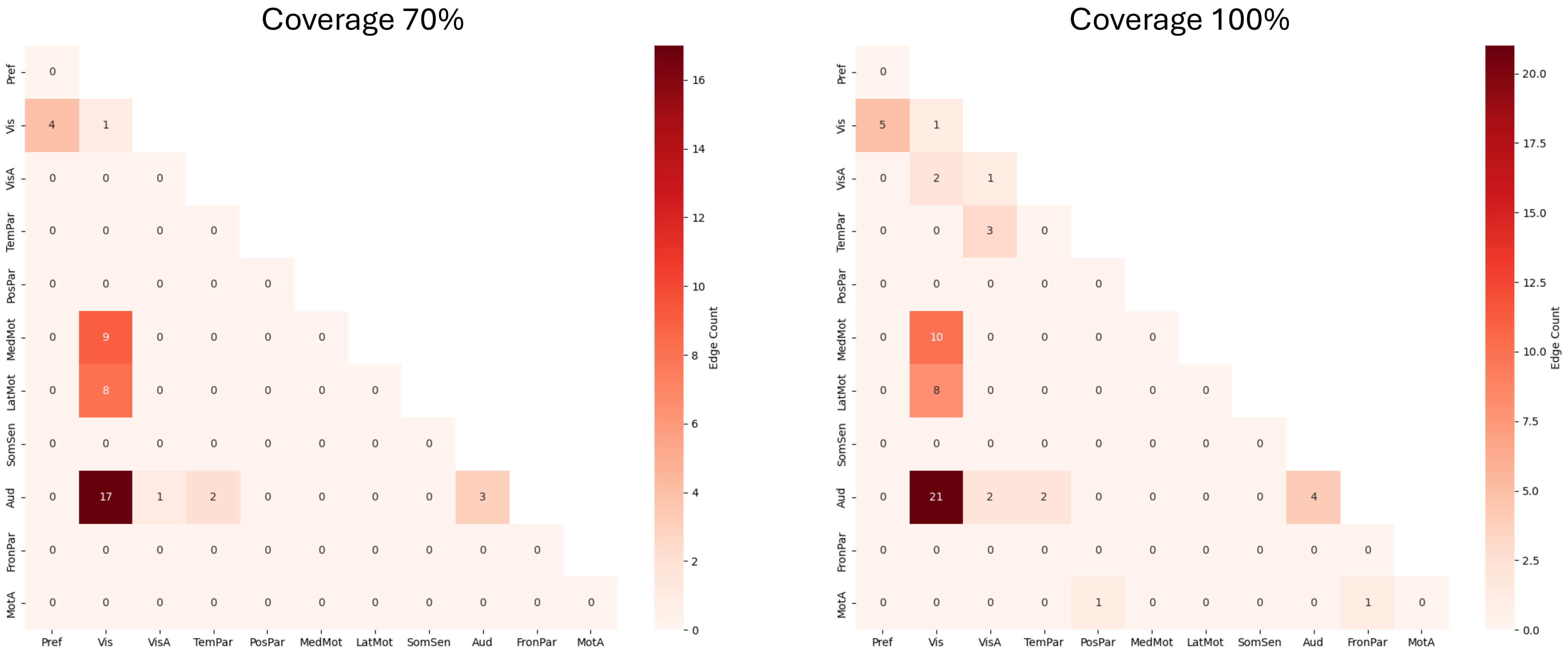


**Figure S14. Comparison between best coverage and full coverage (100%).**
The figure compares best coverage (the coverage level that achieved the highest validation accuracy) with the full coverage condition (100%). The connectivity pattern stays consistent in the two thresholds.

#### Gross motor (whole)

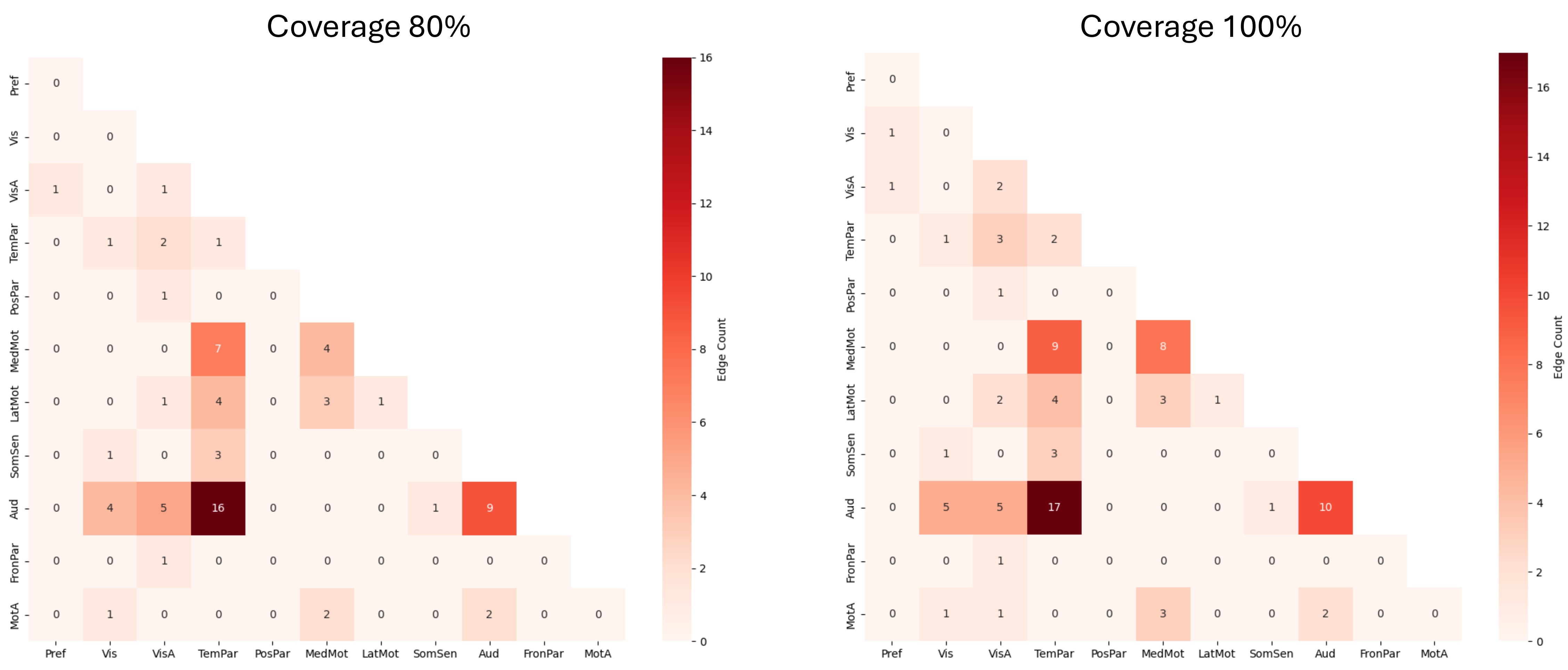


**Figure S15. Comparison between best coverage and full coverage (100%).**
The figure compares best coverage (the coverage level that achieved the highest validation accuracy) with the full coverage condition (100%). The connectivity pattern stays consistent in the two thresholds.

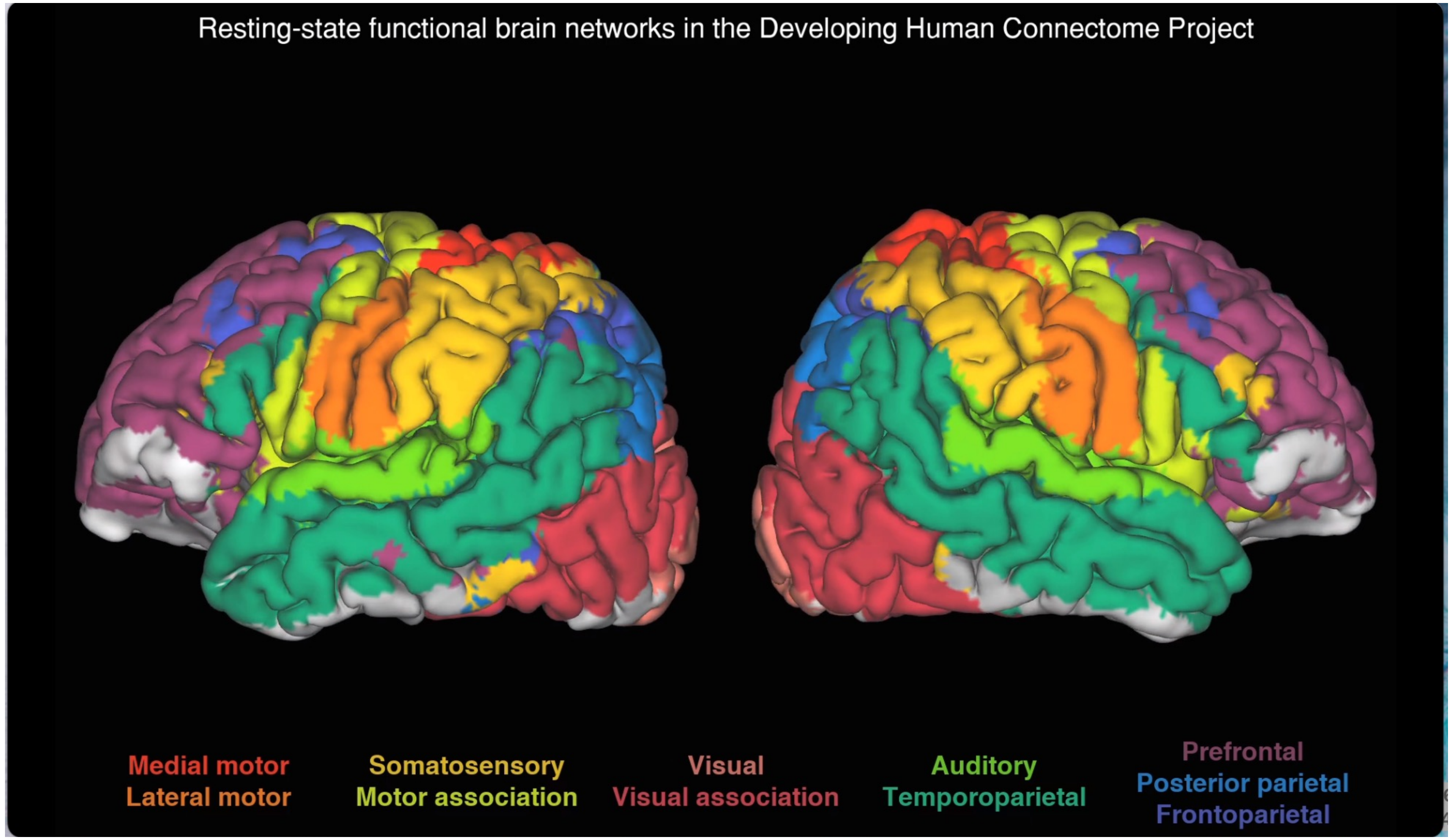


**Figure S16. Network parcellation and RSN assignment.**
Group-level independent component analysis defined canonical resting-state networks (term infants at 43.5–44.5 weeks postmenstrual age).

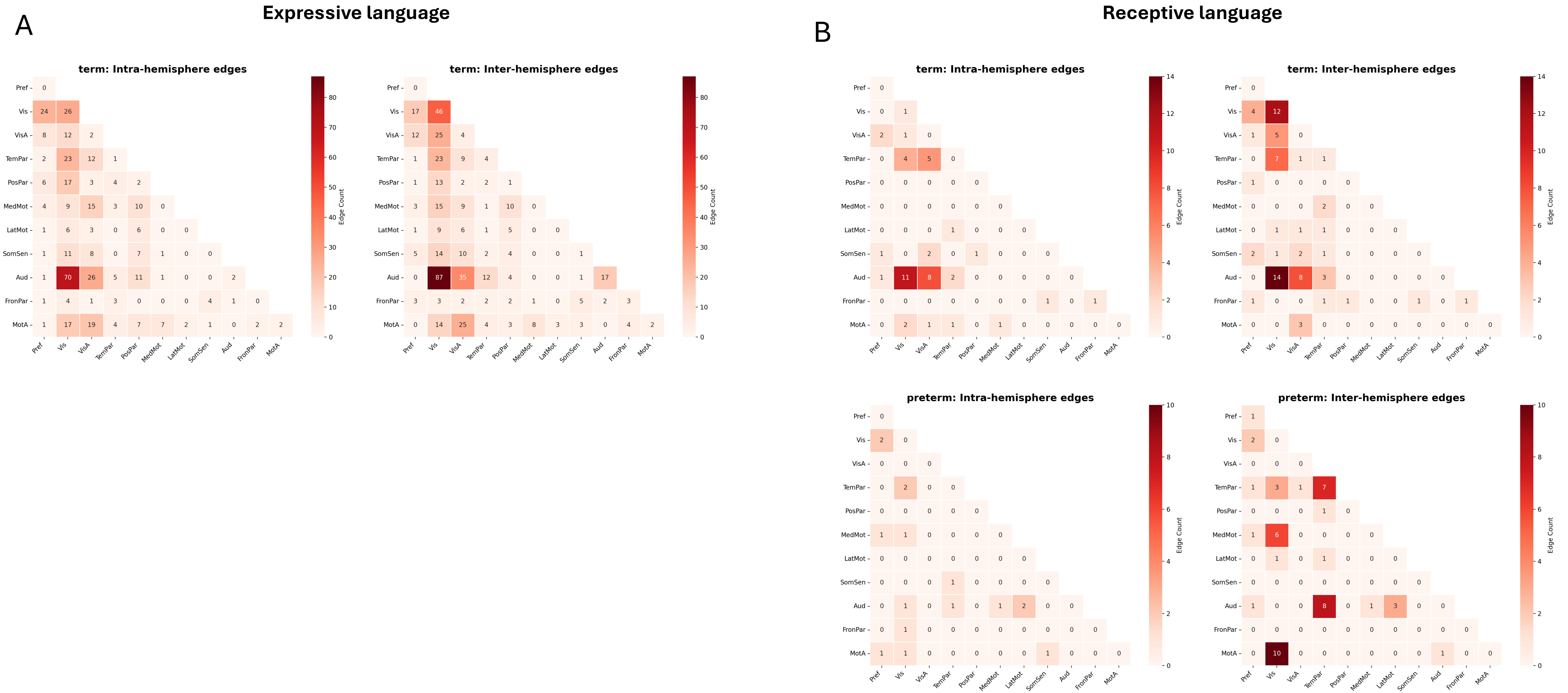
A
Expressive language
term: Intra-hemisphere edges
term: Inter-hemisphere edges
B
Receptive language
term: Intra-hemisphere edges
term: Inter-hemisphere edges
preterm: Intra-hemisphere edges
preterm: Inter-hemisphere edges
Edge Count
Pref
Vis
VisA
TemPar
PosPar
MedMot
LatMot
SomSen
Aud
FronPar
MotA

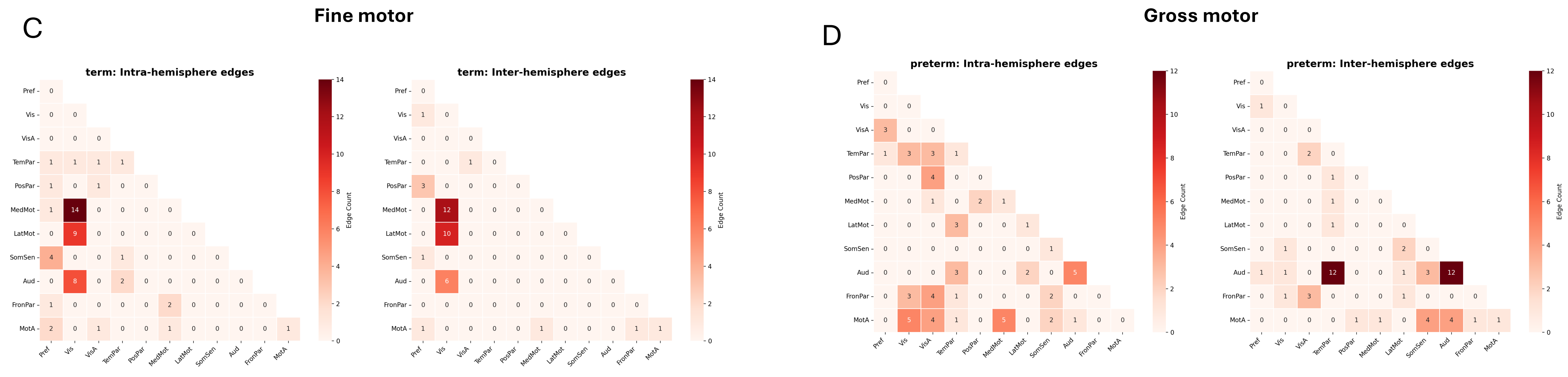


**Figure S17. Inter- vs intra-hemispheric organization differs between term and preterm cohort.** (A,B,C,D) inter and intra-hemisphere between-network connectivity profiles for expressive language, receptive language, fine motor and gross motor respectively.

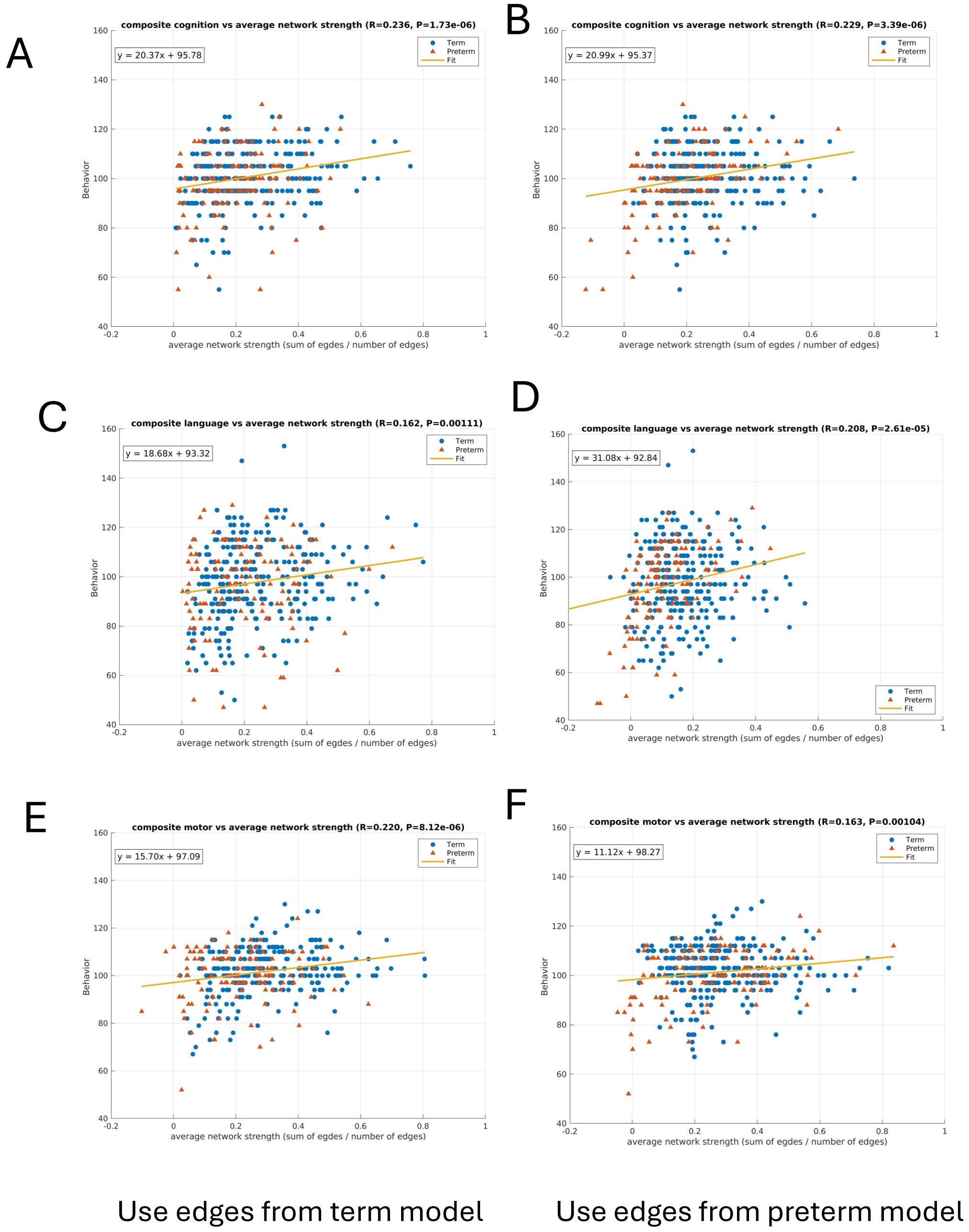


**Figure S18. Average network strength computed using edges derived separately from the term-only and preterm-only models versus behaviour in the whole cohort.**
Scatterplots of average network strength (sum of edges / number of edges) versus Bayley-III composite scores are shown across cognition, language, and motor composite scores (top to bottom). The left panels (A, C, E) use edges derived from the term-only model, and the right panels (B, D, F) use edges derived from the preterm-only model. Behaviour significantly relates to average network strength for both edge sets ($p < 0.001$), indicating the whole-cohort model contains contributions from both term-typical and preterm-typical connectivity patterns.
